# How Filopodia Respond to Calcium in the Absence of a Calcium-binding Structural Protein: They Use Rapid Transit

**DOI:** 10.1101/2021.08.08.455563

**Authors:** C. A. Heckman, O. M. Ademuyiwa, M. L Cayer

**Affiliations:** Department of Biological Sciences, 217 Life Science Building, Bowling Green State University, Bowling Green, OH, USA 43403; Center for Microscopy & Microanalysis, Bowling Green State University, Bowling Green, OH, USA 43403

**Keywords:** cell signaling, rapidly recycling pool, transient receptor potential channel, directional persistence, stromal-interacting molecule, plus-end tracking protein

## Abstract

During directional locomotion, cells must reorient themselves in response to attractive or repulsive cues. Filopodia are narrow actin-based protrusions whose prevalence at the leading edge of a migrating cell is related to the persistence of locomotion. Although there is a marked absence of calcium-binding components in their structure, they responded to store-operated calcium entry (SOCE). Here, we used a two-phase protocol to determine how they responded. In the first phase, extracellular calcium was removed and ER calcium lowered by blocking reuptake through the calcium pump. This was known to activate stromal interacting molecule (STIM) and cause its microtubule-mediated translocation to the cell surface. In the second phase, extracellular calcium and calcium influx into the ER were restored. ER depletion caused filopodia to increase, followed by a spontaneous decrease that was blocked by inhibiting endocytosis. The intracellular calcium concentration increased during depletion, while the size of the exchangeable compartment of vesicles, measured by fluid-phase marker uptake, shrank. When SOCE mediators and the aquaporin, AQP4, were localized, STIM and transient receptor potential canonical (TPRC) channels occupied vesicular profiles side-by-side in linear arrays. STIM1 was translocated, as expected. TRPC1 was initially in a rapidly recycling pool (RRP) where it partially colocalized with Vamp2. Calcium restoration caused TRPC1 exocytosis, while STIM1 reverted toward its original pattern associated with the ER. The exchangeable compartment was restored and this enabled filopodia extension, which was blocked by inhibitors of TRPC1/4/5 and endocytosis. That vesicle recycling was essential for extension during calcium readdition was indicated by reversal of the effect of endocytosis inhibitors in the depletion and readdition phases. The results suggest that SOCE regulates the size of the RRP in epithelial cells, and vesicle recycling is the immediate mechanism affecting filopodia extension. The conclusions are discussed in light of factors regulating protrusion formation, namely surface tension and vesicle trafficking.

## 1 Introduction

Cell polarity mechanisms are essential for the development and differentiation of nearly all tissues. They are especially important in the epithelial lining tissues, where the loss of polarity is recognized by pathologists as one of the key indicators of neoplasia. Studies of cells *in vitro* have shown that the maintenance of polarity relies on the dynamics of filopodia assembly and disassembly. This is also true of cells *in vivo*, particularly the endothelial tip cells, where filopodia at the leading edge increase the efficiency of cell advancement in angiogenic sprouting (Sawamiphak et al., 2010). Filopodia fulfill a similar function in the neuronal growth cone where they mediate guidance, i.e. the directional changes needed for efficient axon navigation through nonneuronal tissues. The receptors for vascular endothelial growth factor (VEGF), VEGFR2, and VEGFR3 are localized on filopodia of the tip cell, and its directional migration is guided by the gradient of VEGF (Gerhardt et al., 2003;Tammela et al., 2008). Likewise, receptors for insulin-like growth factor 1 and epidermal growth factor (EGF) are concentrated on filopodia (Lidke et al., 2005;Krndija and Fairhead, 2019). In general, the persistence of locomotion is closely correlated with the prevalence of filopodia (Heckman et al., 2009). However, the filopodia are sites where signals from substrate adhesion are brought together with cues from chemotactic stimuli. How these signals are interpreted and transmitted to the cell as a whole and how they help determine polarity, remain to be determined.

The processes involved in store-operated calcium have been implicated in polarity determination. SOCE is stimulated by the release of intracellular calcium from the ER downstream of receptor ligation. The Ca^2+^ efflux is triggered by inositol 1,4,5 trisphosphate (IP_3_), which stimulates the opening of a Ca^2+^ channel in the ER called IP_3_ receptor (IP_3_R). The reduced Ca^2+^ concentration in the ER lumen activates the ER Ca^2+^ sensor, stromal interaction molecule (STIM). Membrane vesicles containing STIM associate with adenomatous polyposis coli and end-binding proteins, two types of plus-end, microtubule-associated proteins (Grigoriev et al., 2008;Asanova et al., 2014), and translocate to the plasma membrane where STIM forms a complex with the Orai CRAC (Ca^2+^ release-activated Ca^2+^ channel) transporter. Although Orai1 and STIM are indispensable components of SOCE, TRP channels are often activated downstream resulting in a SOC current, so called because it is induced by SOCE (see for review (Derler et al., 2016)). Thus, TRP channels are thought to amplify the Ca^2+^ signals initiated by STIM and Orai.

Recent studies of neutrophils *in vivo* and *in vitro* provide some insight into how SOCE is involved in locomotion. In neutrophils, the ability to reorient toward a stimulus depended on TRPV (Beerman et al., 2015) or TRPC channels (Lindemann et al., 2013). Because of signaling from filopodia-bound receptors, mentioned above, it is easy to suppose that channels such as Orai and TRP were activated locally near filopodia. However, there is no obvious relationship between Ca^2+^ influx and filopodia dynamics. A structural mechanism, such as that regulating the myosin motor, would depend on a Ca^2+^-binding structural protein. In the sarcomere, this role is filled by calponin C, a protein that changes conformation upon Ca^2+^ binding, resulting in displacement of tropomyosin from the actin filament. In filopodia, a model for activity induced by Ca^2+^ influx, cannot be invoked. The structural constituents of filopodia are well-known, and there is a marked absence of Ca^2+^-binding proteins (Faix and Rottner, 2006;Mattila and Lappalainen, 2008;Faix et al., 2009;Heckman and Plummer, 2013;Jacquemet et al., 2019). A model of Ca^2+^-initiated contraction is therefore irrelevant. Despite the absence of such a structural protein, filopodia respond to Ca^2+^ signals, especially those generated by SOCE. Genetic ablation of the TRPC1 gene prevented the formation of filopodia on the tip cells during sprouting angiogenesis (Yu et al., 2010). Both STIM and TRPC were essential for axon guidance, and the attractive turning of the growth cone could be switched to repulsive turning by knock-down of either protein (Wang and Poo, 2005;Mitchell et al., 2012;Shim et al., 2013).

Both STIM and TRPC are implicated in the determination of polarity via the highly conserved mechanism for maintaining a cell’s anterior-posterior axis, phosphoinositide 3-kinase (PI3-K) and PTEN (Phosphatase and TENsin homolog deleted on chromosome 10). Patterns of TRPC and STIM trafficking were subject to regulation by PI3-K and PTEN, suggesting that SOCE participated in regulating this axis (Kini et al., 2010;Monet et al., 2012;Chaudhuri et al., 2016). These considerations are complicated by the complexity of Ca^2+^ signaling, because its concentration in cells undergoes dynamic fluctuations, which occur on both a global and a local scale. In migrating cells, the Ca^2+^ concentration is low at the leading edge and high at the rear of the cell. Within the lowest portion of this gradient, calcium flickers were found (Taylor et al., 1980), which are sites where the IP_3_R is activated (Wei et al., 2009). Ca^2+^ influx at the leading edge was implicated in sustaining PI3-K activity and ruffling, which in turn drive forward movement of the cell (Evans and Falke, 2007).

Filopodia exhibit repetitive cycles of extension, stabilization, and retraction, often extending repeatedly from the same sites over several cycles of extension and retraction (Schäfer et al., 2011). Where focal adhesions were found in the lamellipodium interior to such staging areas (Nemethova et al., 2008;Shutova et al., 2012), they had a longer lifetime than in areas lacking filopodia (Schäfer et al., 2009). The cycles of extension and retraction depend on tension on these adhesions, which may in turn be regulated by feedback loops between activated Cdc42 and Rac (Schäfer et al., 2010;Schäfer et al., 2011). Myosin II maintains tension on the actin bundle, and some evidence suggests that filopodia lifetime depends on its pulling force (Alieva et al., 2019). Adhesive sites containing vinculin recruited myosin X, which is implicated in filopodia initiation (He et al., 2017). The importance of tension is emphasized by additional findings, first, that the tension force exerted by filopodia was greater on stiffer substrates (Athamneh and Suter, 2015). Secondly, the prevalence of filopodia was higher on substrates of greater adhesiveness (Amarachintha et al., 2015).

Tension has another role in protrusion regulation, however. Centripetal forces on the plasma membrane, maintained by the pushing force of actin on the membrane and by in-plane tension of the lipid bilayer, limit the cell’s ability to form protrusions. This must be overcome by an opposing force, which is estimated at 60 pN for the filopodium and is generated by the polymerization of the actin filaments making up its structural core (see for reviews (Heckman and Plummer, 2013;Lieber et al., 2013;Athamneh and Suter, 2015;Verkhovsky, 2015)). The centrifugal force may facilitate positive feedback. As it is developed at the tip of a filopodium and transmitted to the base, it may contribute to the tension on adhesions in the adjacent lamellipodium (see for review (Jacquemet et al., 2015)). Another means of regulating filopodial extension and retraction was proposed recently, namely supplying or withdrawing membrane at the sites where filopodia are initiated (see for review, (Gallop, 2020)). This hypothesis holds that essential membrane components could be deposited or withdrawn depending on the exo- and endocytic activities taking place during initiation and extension. While these concepts are not mutually exclusive, each may apply to only one phase of the dynamic cycle, namely initiation, extension, stabilization, or retraction. The dependence of filopodia on SOCE could involve any one of these phases.

In the current research, we removed exogenous stimuli and determined how intracellular calcium levels, [Ca^2+^]_i_, were regulated by SOCE initiation. The Ca^2+^ concentration in the ER is many-fold higher than in the cytoplasm but is lowered by Ca^2+^ efflux through IP_3_R, as illustrated in Supplementary Figure 1A, thus initiating SOCE. In the current experiments, the SOCE process was broken down into separate phases of ER depletion and readdition of Ca^2+^. The results show how the distribution of STIM, Orai, and TRPC1 were changed and contrast their redistribution with that of AQP4, an aquaporin in the respiratory airway lining cells. The location of a TRP channel, TRPC1, which was closely related to filopodia dynamics, was regulated by trafficking to and from the rapidly recycling pool (RRP) of vesicles.

## 2 Methods

### 2.1 Cell culture and treatment with pharmacological agents

An established rat cell line, 1000 W, was used. It was originally generated from rat tracheal epithelium as a model of human bronchogenic carcinoma (Marchok et al., 1978). The cells were maintained in a modified Waymouth’s medium (Sigma-Aldrich, St. Louis, MO) containing penicillin, streptomycin, 10% fetal bovine serum (Hyclone, UT or Atlanta Biologicals, GA), 0.1 µg/ml insulin, and 0.1 µg/ml hydrocortisone, as previously described (Heckman et al., 2017). They were subcultured by detachment with a trypsin solution (Invitrogen, Grand Island, NY) made up in Ca^2+^-, Mg^2+^-free Hanks’ balanced salt solution (Ca^2+^-free HBSS, GIBCO, Gaithersburg, MD).

To inhibit TRPC and CRAC channels, SKF963665 (1-(beta-[3-(4-methoxy-phenyl)propoxy]-4-methoxyphenethyl)-1H-imidazole hydrochloride) was obtained from Selleckchem (Munich, Germany). SKF96365 blocked nonselective cation channel activity with IC50s of 3 to 16 µM (Merritt et al., 1990;He et al., 2005). Pico145 (Chem Scene, Monmouth Junction, NJ) was used at 0.3-1.3 nM to inhibit TRPC1/4/5 isoforms. Nifedipine (MedChem Express, Monmouth Junction, NJ) was used to inhibit voltage-activated calcium channels (VACC). The reagents were used at final concentrations of 1-3 times the IC50 for the respective activities (Merritt et al., 1990;Schrøder et al., 2008), except for CRAC and VACC channel inhibitors. Although these compounds were active against the respective channels at micromolar concentrations (Franckowiak et al., 1985;Zitt et al., 2004;Wang et al., 2018), they had no effect on filopodia at these concentrations and so higher concentrations were used.

Possible mediators of the response to altered [Ca^2+^]_i_ were investigated by using cell-permeable peptides, enzyme inhibitors, and receptor agonists. 1,2-dioctanoylglycerol (DOG) was obtained from Sigma-Aldrich, and the aldehyde inhibitor of calpains, ALLN, from Focus Biomolecules (Plymouth, PA). Calcium-like peptide 2 (CALP2) and N-(6-aminohexyl)-5-chloro-1-naphthalenesulfonamide hydrochloride (W-7) were obtained from Tocris (Minneapolis, MN). CaM kinase II and calcineurin inhibitors, autocamtide-2-related inhibitory peptide and calcineurin autoinhibitory peptide were obtained from EMD Millipore, Temecula, CA. Myosin light chain kinase (MLCK) inhibitors, ML-7 and MLCK peptide, were obtained from Cayman Chemicals (Ann Arbor, MI). When epidermal growth factor (EGF) was used to supplement Ca^2+^-free HBSS, the human recombinant protein (PeproTech, Rocky Hill, NJ) was used at a final concentration of 10 ng/ml.

### 2.2 Sample preparation, filopodia counts, and statistical analysis

Cells were fixed with warm, buffered 3% formaldehyde (pH 7.4) made fresh from paraformaldehyde in cytoskeletal buffer. The samples were rinsed with phosphate-buffered saline and stored in buffer at 4° C until observations were made. Samples were mounted on slides and assigned code numbers before being examined, and counts were made by independent observers who had no knowledge of the sample’s identity. Filopodia prevalence was determined on single cells by analyzing the fraction of cells with filopodia and the proportion of their perimeter covered with filopodia. Filopodia morphology is also that previously described (Heckman et al., 2017). Because the coverage of the cell perimeter varied from one experiment to another, counts are presented relative to counts in the sham-treated control. Where experiments were concluded by fixation after Ca^2+^ readdition, counts were indexed to the counts after ER depletion, i.e. the starting point for filopodia extension. Microsoft Excel was used to calculate averages and standard deviations.

An online service, https://www.statskingdom.com/320ShapiroWilk.html, was used to test the filopodia control sample means of 20 experiments. A probability value of P= 0.342 was obtained, indicating a low probability that the populations deviated from the normal distribution. Statistical differences among the means of experimental values were evaluated using the online service for one-way ANOVA, https://www.socscistatistics.com/tests/anova/default2.aspx, and the Tukey test was used for multiple comparisons. Comparisons between individual treatment groups and controls were done by the two-tailed Student’s t-test with Bonferroni correction for multiple comparisons. All error bars shown represent ± one standard error of the mean (S.E.M.)

### 2.3 Germanium substrates for cell culture

To make the substrates adhesive for 1000W cells, a film of germanium (Structure Probe, Inc., West Chester, PA) was applied to glass coverslips of 25 mm diameter and thickness #1 (Electron Microscopy Sciences, Hatfield, PA) as previously described (Heckman et al., 2017). Coverslips were sterilized by ultraviolet irradiation and placed in 35-mm culture dishes. For experiments, 2–3×10^5^ cells were plated in each 35-mm dish and left overnight to become attached.

### 2.4 Calcium store depletion and replenishment

To deplete the ER calcium store, the culture medium was replaced with Ca^2+^-free HBSS containing

1.5 µM ethylene glycol-bis(β-aminoethyl ether)-N,N,N’,N’-tetraacetic acid (EGTA) or 5 µM cyclopiazonic acid (CPA, CalBiochem-EMD Millipore). For Ca^2+^ replenishment, the coverslips with the cells attached were transferred sequentially into Ca^2+^-free HBSS for CPA washout (5 minute) and then Ca^2+^-replete Hanks balanced salt solution (HBSS) for variable lengths of time. Although the effects of CPA and EGTA on filopodia were indistinguishable during store depletion, recovery was less reproducible with EGTA. Therefore, CPA was used in the experiments.

### 2.5 Changes in Ca^2+^ concentration

Intracellular calcium levels were compared in cells incubated in Ca^2+^-free HBSS before ER stress and during depletion of the ER with CPA. This was done by exposing cells to calcium orange AM (Invitrogen, Eugene, OR) for 20 minutes in culture medium, then transferring the cells to Ca^2+^-free HBSS with or without CPA. Calcium orange was made up at 0.25 µg/µl in dimethylsulfoxide and used at a final concentration of 2 µg/ml. Images were acquired at 5-minute intervals, using the software, instrumentation and settings described below (see **2.8 Equipment and settings**) and processed by background subtraction. The averaged intensity value per cell was calculated for each frame processed.

### 2.6 Assay for horseradish peroxidase (HRP)

To measure fluid-phase uptake, we added an aliquot of stock HRP to cells in either culture medium or in Ca^2+^-free HBSS containing 5 *μ*M CPA. After 30 minutes of treatment, the cells were lysed in 0.1 % Triton X-100 and recovered as described previously (Heckman et al., 1996). The assay mixture was composed by making up aliquots of cell lysate with phosphate buffer (pH 5.0) to a final concentration of 0.05 M. H_2_O_2_ and o-dianisidine were added to final concentrations of 0.003% (v/v) and 80 µM respectively. The reaction rate was measured by following the change in absorbance at 410 nm in a Thermo Scientific Genesys 10 UV spectrophotometer.

### 2.7 Immunofluorescence localization, image acquisition, and image processing

For immunofluorescence staining, cells were fixed as above and permeabilized with 50 µg/ml digitonin (LC Laboratories, Woburn, MA) and 0.2% Triton-X 100 in cytoskeletal buffer. Mouse monoclonal anti-β-tubulin and rabbit polyclonal anti-TRPC1 were obtained from Sigma-Aldrich. Monoclonal antibodies against Vamp2 (R&D Systems, Minneapolis, MN), CaV1.2 (Novus Biologicals, Centennial, CO), caveolin-2 (Thermo Fisher, Rockford, IL), and STIM1/CRACR2A (CRAC regulator 2A, Cedarlane Laboratories, Burlington NC) and polyclonal antibodies against Orai1 and Orai3 (ProSci, Poway, CA) were made up in cytoskeletal buffer according to the manufacturers’ specifications. Rabbit antibody against aquaporin 4 (AQP4), which is present in most cell types of the bronchus and trachea, was a gift of Søren Nielsen (Nielsen et al., 1997). AQP3, also present in the airway, could not be identified in 1000W cells by immunolocalization (data not shown).

In experiments where sheep antibody against the extracellular loop of TRPC1 (Antibodies-online, Limerick, PA) was used with anti-AQP4 or anti-caveolin, primary staining was followed by Cyanine 3-labeled donkey anti-sheep and a fluorescein isothiocyanate (FITC)-conjugated antibody against either rabbit or mouse antibody (Jackson ImmunoResearch, West Grove, PA). Otherwise, secondary staining was performed using donkey anti-mouse tetra-rhodamine-labeled and goat anti-rabbit FITC-labeled antibodies (Jackson ImmunoResearch). Triple staining procedures were done using the secondary antibodies, donkey anti-mouse (Abcam, Cambridge, MA) tagged with Cyanine5, Cyanine 3-tagged anti-sheep, and FITC-tagged anti-rabbit. Samples were mounted in 2.5% DABCO made up in 2,2′-thiodiethanol (Sigma-Aldrich) and viewed with a 100x lens as described below (see 2.8 **Equipment and settings**). Control samples exposed to irrelevant primary antibodies, followed by the same secondary antibodies as above, showed no staining.

### 2.8 Equipment and settings

A confocal Leica DMI3000B inverted microscope (Leica Microsystems, Buffalo Grove, IL) equipped with a Lumen Dynamics X-Cite light engine and Leica rhodamine filter set (Excitation 546/10 nm, dichroic LP 560 nm, Emission 585/40 nm) and 10x lens were used to assess intracellular Ca^2+^ concentration. Images were acquired once per minute with a Rolera Thunder cooled CCD camera with back-thinned, back-illuminated, electron-multiplying sensor (QImaging, Surrey, British Columbia, Canada). MetaMorph version 7.8, 4.0, (Molecular Devices, Sunnyvale, CA) was used to configure the hardware settings for time-lapse recording.

Samples triple-stained for the SOCE mediators, STIM1, Orai1/3, and TRPC1, were also imaged with the Leica DMI3000B using Spectra X LED source (Lumencor, Beaverton, OR) and X-Light spinning-disk confocal unit (CrestOptics, Rome, Italy) with Semrock bandpass filter FF01-440/521/607/700-25 and dichroic FF410/504/582/669-Di0l-25×36. Software and CCD camera were those identified above.

For studies by higher magnification, epifluorescence images were taken on an Axiophot microscope (Carl Zeiss, Jena) using Zeiss 100x Plan-Neofluar/1.30 objective lens and FluoArc mercury lamp. Filter sets for FITC (excitation 490 nm, emission 525 nm) and Cyanine3 or tetrarhodamine (excitation 545 nm, emission 605 nm) were from Chroma (Bellows Falls, Vermont). Images were acquired with an Andor camera (Zyla 4.2 PLUS sCMOS, Concord, MA) running under Molecular Devices MetaVue software. The final pixel size was 63 nm.

### 2.9 Refractive index determination

To determine the role of water entry in filopodia formation, bovine serum albumin (Fisher Fraction V, Heat Shock Treated) was made up in Waymouth’s medium (see **2.1 Cell culture and treatment with pharmacological agents)** at 33.35% (w/v) and diluted 3:4 with medium before use. Cells were cultured on germanium coverslips and maintained at 37 °C while being viewed. Time-lapse recordings were made with a phase lens (Plan-apochromatic 100x) in the Zeiss Axiophot, with image acquisition as described above (see **2.8 Equipment and settings)**.

### 2.10 Correlation coefficients

Pearson correlation coefficients were obtained for matched images of thin, peripheral areas viewed in the FITC and tetra-rhodamine channels. The Colocalization Finder (Laummonerie and Mutterer, 2004) plugin, running under ImageJ (Schneider et al., 2012), was used. With this plugin, background outside the cells and thicker parts of the interior could be excluded from analysis. Because we did not attempt to remove the background, the coefficients included fluorescence background surrounding the particles. This was nonspecific, so a value around 0.4 was the lowest average coefficient observed.

### 2.11 Analysis of particles’ dimensions after staining by indirect immunocytochemistry

The different loci, represented by immunocytochemical localizations of their intracellular sites, were segmented from images of thin, peripheral portions of the cells. The selection of these areas obviated the complication of multiple layers of structures, as overlapping layers of particles were absent. Contrast enhancement and edge detection operations were run on each image in ImageJ, then the threshold was tuned to maximize the number of complete, continuous outlines around figures. The size of the figures was measured in pixels, using the particle analysis plugin and a circularity setting of 0.9-1.0. The radius was calculated by modeling the number of pixels in each locus as a circle, and the relative size distributions compared on plots.

To determine the diameter of the zone occupied by TRPC1 clusters, called particles, at the cell surface, we estimated the maximum extent of the zone around each filopodium using the line function of ImageJ. All particles within the zone of each filopodium were assigned to the maximum distance, and the results were plotted as a cumulative distribution of distances.

## 3 Results

### 3.1 Changes in filopodia prevalence during the SOCE procedure

A classical two-step procedure was used to separate the SOCE process into two processes (Putney, 1991). First, cells were deprived of Ca^2+^ in the presence of CPA to deplete the ER of Ca^2+^ and form the STIM1-Orai complex (see for reviews (Carrasco and Meyer, 2011;Soboloff et al., 2012)). In the second step, CPA was washed out and the Ca^2+^-free medium was replaced with Ca^2+^-replete medium to initiate refilling of the ER with Ca^2+^. This procedure caused complex changes in filopodia formation. Ca^2+^ depletion caused a transient increase followed by a spontaneous decline to the starting level by 30 minutes (Figure 1A and B). Upon Ca^2+^ replenishment, filopodia increased again for up to 20 minutes.

**Figure 1.**
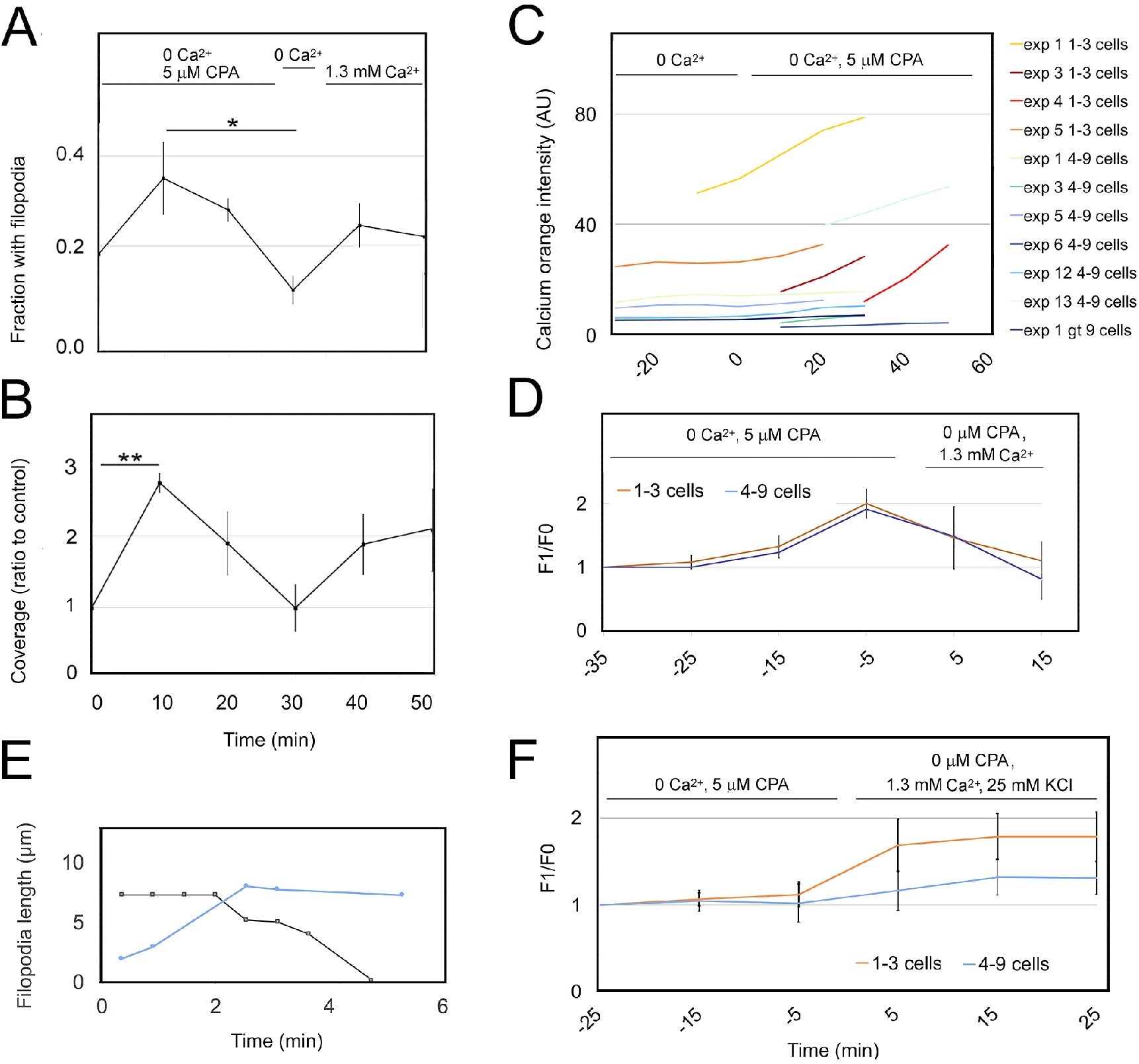
Induction of SOCE causes an increase in [Ca^2+^]_i_ and transient changes in filopodia. Cells were exposed to 5 *μ*M CPA or 1.5 µM EGTA in Ca^2+^-free HBSS to deplete the ER of Ca^2+^. Results in panels A-B and D-F represent 3-7 experiments. (A, B) Ca^2+^ replenishment was initiated by CPA washout and replacement of extracellular Ca^2+^. At various times during ER depletion and replenishment, samples were fixed and the fraction of cells with filopodia and percent of the perimeter covered with filopodia were determined. (A) Significance by ANOVA, P=0.0017. *Treatments differ at P=0.0015. (B) Significance by ANOVA, P=0.0030. **Treatments differ at P=0.0028. (C) Fluorescence intensity of groups of cells preloaded with calcium orange and then exposed to Ca^2+^-free HBSS with or without CPA. CPA was added at time=0. The intensity is shown as the moving average of two time points to eliminate short-term fluctuations. Areas with 1-3 cells (yellow and red) and colonies of cells (blue and green) were analyzed separately. Significance of experiment by ANOVA repeated measures, P<0.0001. (D) Fluorescence intensity of calcium orange during ER depletion and replenishment of extracellular Ca^2+^. Depletion is represented by negative numbers and replenishment by positive numbers, and emission is shown as the ratio of the intensity over initial intensity (F_1_/F_0_). The means, tested by ANOVA repeated measures, differed significantly at P=0.0042. (E) Filopodia extension and retraction. Typical extension (blue) and retraction (black) rates are 1.7 µm/minute and 1.4 µm/minute, respectively. The mean length was 5.7 µm (S.D. 1.0 µm, average from 8 images). (F) Increase in F_1_/F_0_ after CPA washout followed by Ca^2+^ with 25 mM KCl, which was supplied at time=0. Values after time=0 differed significantly by the ANOVA repeated measures test, P<0.0001. Error bars represent ± one standard error of the mean (S.E.M.).

#### 3.1.1 Ca^2+^ depletion in the ER causes a sustained rise in [Ca^2+^]_i_ and transient filopodia formation

To determine how filopodia formation was related to [Ca^2+^]_i_, we preloaded cells with a Ca^2+^ indicator and visualized the Ca^2+^-dependent fluorescent emission by confocal microscopy. The emission changed little over 30 minutes when the medium was replaced with Ca^2+^, Mg^2+^-free balanced salt solution, consistent with previous results showing that Ca^2+^ deprivation in serum-free medium had little effect on intracellular Ca^2+^ concentration (Davenport and Kater, 1992). Subsequent addition of CPA caused a continuous rise in [Ca^2+^]_i_ for over 40 minutes (Figure 1C). This result was consistent with the expectation that the sarcoplasmic reticulum Ca^2+^-ATPase pump was inhibited, and thus Ca^2+^ could not be pumped back into the ER. Filopodia were inhibited when the [Ca^2+^]_i_ increase continued for over 10 minutes (Figure 1A-C).

[Ca^2+^]_i_ decreased after CPA washout and restoration of extracellular Ca^2+^ (Figure 1D). Again, this was consistent with expectations, because CPA removal would relieve inhibition of the sarcoplasmic reticulum Ca^2+^-ATPase pump and allow Ca^2+^ to be pumped back into the ER store (see for review (Putney and Parekh, 2005)). During this part of the Ca^2+^ depletion-readdition protocol, filopodia increased again for up to 20 minutes, while [Ca^2+^]_i_ decreased to the baseline level of untreated cells (cf. Figures 1A-B and D). The comparison of declining [Ca^2+^]_i_ to filopodia length suggested that filopodia extended at a rate of 1.7 µm/minutes during Ca^2+^ replenishment, and then remained stable for variable lengths of time. They retracted at a similar rate, i.e. 1.4 µm/minutes (Figure 1E). The rates were in the range similar to those of human keratinocytes (Schäfer et al., 2011) and cells from other animal cell lines (for review, see (Bornschlögl, 2013;Heckman and Plummer, 2013)).

Filopodia are well-known to exhibit repetitive cycles of extension and retraction as mentioned above (see **1 Introduction**). To determine how many cycles could take place during the time when filopodia were decreasing, we estimated the lifetime of the filopodia from time-lapse recordings. It was 8.5 minutes (S.D. 6.6, number of filopodia=10), suggesting that the increase in filopodia at 10 minutes after CPA treatment represented one cycle, and filopodia formation was inhibited when 2-3 cycles had elapsed. The effect of depolarization on filopodia prevalence was tested by including 25 mM KCl in the solution used for Ca^2+^ replenishment. As expected, this led to further [Ca^2+^]_i_ elevation compared to restoration of extracellular Ca^2+^ alone (cf. Figures 1D and F), but the filopodia disappeared (see **3.2.3 Exocytosis and stimulus-coupled secretion during extracellular Ca**^**2+**^ **replenishment**).

#### 3.1.2 Filopodia responses depend on store-operated Ca^2+^

During ER depletion, cells are deprived of growth factors from the medium by replacement of the culture medium with a Ca^2+^-free balanced salt solution. This prevented phospholipase C activation and downstream signaling to IP_3_R through IP_3_-mediated calcium release, as shown schematically in Supplementary Figure 1A. To determine whether the deprivation of signaling input caused net filopodia extension and retraction, we assayed the prevalence of filopodia after incubating cell samples with HBSS with or without Ca^2+^. Only trivial changes occurred, compared to those caused by CPA (cf. Figures 1A-B and Supplementary Figure 1B-C).

The induction of Ca^2+^ influx downstream of receptor signaling was a hallmark of SOCE (Putney and Tomita, 2012;Prakriya and Lewis, 2015;Bird and Putney Jr., 2018). To confirm that SOCE was mediating filopodia formation, we sustained phospholipase C signaling to the IP_3_R, by providing EGF during the first part of the Ca^2+^ depletion-readdition protocol (see **3.1 Changes in filopodia prevalence during the SOCE procedure**) and measuring filopodia prevalence after Ca^2+^ influx into the ER was restored. When extracellular Ca^2+^ was replenished after 20 minutes of ER depletion in the absence of EGF, there was little net extension. In contrast, signaling and IP_3_ production continued in cells treated with EGF and CPA, driving STIM1 activation and SOCE induction, and filopodia recovered to the same extent as with a 30-minute interval of depletion (cf. Figure 1A-B and Supplementary Figure 1D-E). The results were consistent with the expectation that IP_3_ signaling increased Ca^2+^ efflux from the ER and hence the driving force for SOCE. In certain cell types, the ryanodine receptor released Ca^2+^ in response to increases in [Ca^2+^]_i_. To determine whether ryanodine receptors played a role in filopodia formation, we treated cells by Ca^2+^ depletion in the presence or absence of the receptor agonist, caffeine. When the filopodia were analyzed after restoration of extracellular Ca^2+^, their prevalence would have been greater in caffeine-treated cells due to enhanced Ca^2+^ efflux from the ER, if ryanodine receptors were present. This had little effect (Supplementary Figure 1F-G), indicating that these receptors contributed little to ER depletion.

#### 3.1.3 A calmodulin inhibitor rescues filopodia during ER depletion

Although the lifetime of the filopodia was variable, it was sufficient to allow 2-3 cycles of retraction and extension after the net formation following Ca^2+^ removal (see **3.1.1 Ca**^**2+**^ **depletion in the ER causes a sustained rise in [Ca**^**2+**^]_**i**_ **and transient filopodia formation**). It could be argued that filopodia formation was blocked because a calcium-activated protein caused the disappearance of an essential molecule from the plasma membrane. Previous work implicated calpain, along with two Ca^2+^/calmodulin-activated enzymes, calcineurin and CaMKII, in filopodia retraction and extension in the growth cone (Kerstein et al., 2017) (Tojima et al., 2011). When cells were treated with inhibitors of CaMKII, calcineurin, calpain, and Ca^2+^/calmodulin, however, only the calpain inhibitor, ALLN, and the Ca^2+^/calmodulin antagonist, CALP2, rescued filopodia (Figure 2A-B). ALLN had the opposite effect in the growth cone, where the calpain inhibitor had decreased the average filopodial lifetime and reduced the number of stable filopodia (Robles et al., 2003). Not only was the result opposite, but the higher concentration of 250 µM caused the cells to round up. This meant the majority of the cell edge was not visible precluded further analysis of its effects. CALP2 caused a large increase in filopodia, which was suggestive in light of the fact that the concentration of Ca^2+^/calmodulin might increase due to the persistent increase in [Ca^2+^]_i_. Intermediaries in signaling upstream of STIM or a blockade in the rise in [Ca^2+^]_i_ were considered unlikely to mediate the effect of CALP2, because [Ca^2+^]_i_ in upper airway epithelial cells had been altered very little by CALP2 treatment (Ten Broeke et al., 2001).

**Figure 2.**
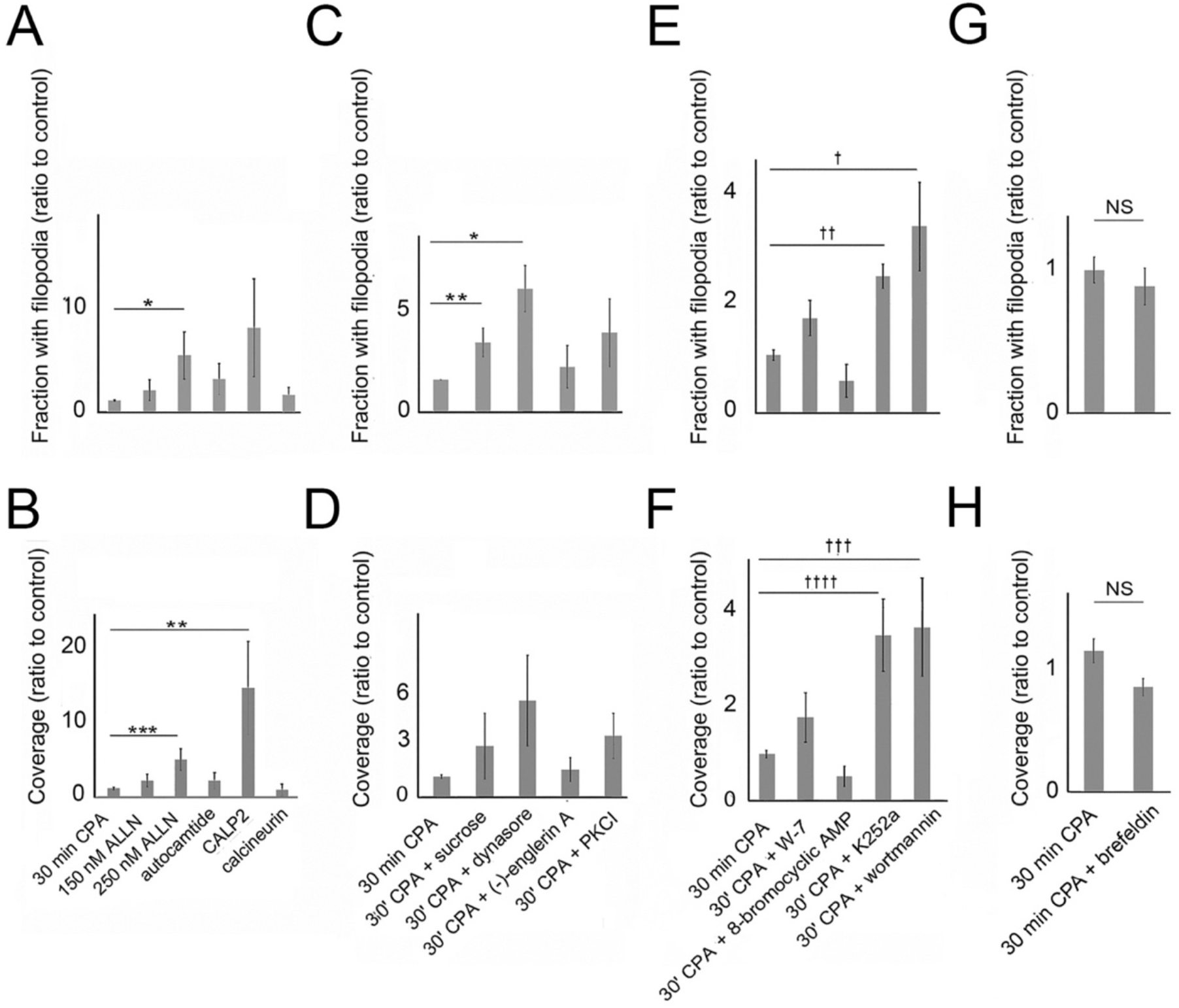
Tests of possible effectors mediating filopodia disappearance during ER depletion. Cells were exposed to 5 *μ*M CPA in Ca^2+^-free HBSS to deplete the ER of Ca^2+^. Inhibitors were used at final concentrations of 1-3 times the IC50 for the targeted activity. Means of 3-7 experiments are shown ± S.E.M. (A, B) Filopodia prevalence in the presence or absence of an inhibitor of calpain (ALLN), CaMKII (10 µM autocamtide-2 related inhibitory peptide), Ca^2+^/calmodulin (20 µM CALP2), or calcineurin (20 µM calcineurin-autoinhibitory peptide II). (A) Significance by ANOVA, P=0.035. *Treatments differ at P=0.037. (B) Significance by ANOVA, P=0.0089. **P=0.038. ***P=0.0045. (C, D) Filopodia prevalence in the presence or absence of 0.4 M sucrose, 48 µM dynasore, 23 nM (-)-englerin, or 24 µM protein kinase C inhibitor (PKCI). (C) Significance by ANOVA, P=0.0023. *Treatments differ at P=0.0032. **P=0.041. (D) Significance by ANOVA, P=0.0032. (E, F) Filopodia prevalence in the presence or absence of W-7 (60 µM), 8-bromocyclic AMP (80 µM), K252a (50 nM), or wortmannin (1 µM). (E) Significance by ANOVA, P=0.00001. ^†^Treatments differ at P=0.0032. ^††^P=0.0004. (F) Significance by ANOVA, P=0.00001. ^†††^Treatments differ at P=0.0056. ^††††^P=0.0012. (G, H) Filopodia prevalence in the presence or absence of 20 µg/ml brefeldin A.

In neurons, calmodulin is a Ca^2+^ sensor and regulates endocytosis (see for review (Wu et al., 2014)). The possibility that CALP2 inhibited endocytosis was tested by comparing its effect with that of dynasore, an inhibitor of clathrin-mediated endocytosis. Dynasore enhanced filopodia while showing less variability. Other inhibitors of endocytosis, i.e. hypertonic sucrose and a myristoylated pseudosubstrate sequence of protein kinase C (PKC) α/β, PKCI, had similar effects, albeit weaker (Figure 2C-D). A target of the Ca^2+^/calmodulin effect on endocytosis is not known, so the CALP2 effect may rely on multiple Ca^2+^/calmodulin-regulated activities among the hundreds of proteins potentially regulated, as mentioned above. Ubiquitous downstream calmodulin targets, phosphodiesterase and MLCK, were ruled out as mediators by pharmacological experiments. Phosphodiesterase activation would be counteracted by a calmodulin antagonist such as CALP2, and this would allow the accumulation of cAMP. Treating cells with the cAMP analogue, 8-bromocyclic AMP, had an effect opposite that predicted (cf. Figure 2A-B and E-F). Likewise, treatment with MLCK inhibitors indicated that this enzyme was unlikely to mediate the effect of Ca^2+^/calmodulin. MLCK was essential for the cyclical protrusion and retraction of the lamellipodium and filopodia extension in fibroblasts. In longer-term exposures, MLCK inhibitors, ML-7 (Hu et al., 2014) and altenusin (Varghese et al., 2012) decreased filopodia. Another way in which MLCK could affect filopodia was by its promotion of plasma membrane TRPC5 channel distribution and channel activity (Shimizu et al., 2006). All of these activities suggested MLCK would enhance filopodia after an increase in Ca^2+^/calmodulin, and MLCK inhibitors would be inhibitory. This was not observed. MLCK inhibitors, altenusin, MLCK peptide, and ML-7, in combination with CPA had little effect (data not shown). K252a is a broad-spectrum kinase inhibitor, affecting both MLCK and protein kinase A at IC50s of 10-50 nM. It was expected to decrease filopodia rather than increase them, if working through its effect on MLCK, but K252a had the opposite effect. K252a may have increased filopodia by inhibiting protein kinase A, as this would be consistent with its activator, 8-bromocyclic cAMP, having the opposite effect to that of K252a. Not only was the inhibition of protein kinase A one possible explanation of the K252a effect, but K252a also inhibits PKC. This would replicate the effect of PKCI, and the combined effect on protein kinase A and PKC together could account for the observations with K252a (cf. Figure 2C-D and E-F).

Among the numerous proteins potentially affected by calmodulin, there were many channels that respond to Ca^2+^/calmodulin-dependent inactivation (see Supplementary Material, **Targets of Ca**^**2+**^**/calmodulin**). If Orai or TRPC functions were ordinarily inhibited by Ca^2+^-dependent inactivation, but rescued by CALP2, it would have enhanced Ca^2+^ influx under normal conditions (Singh et al., 2002;Litjens et al., 2004;Mullins et al., 2009), but not under the conditions of the current experiments, where extracellular Ca^2+^ was absent. To test whether hypothetical Ca^2+^-dependent inactivation of TRPC channels could account for the inhibition of filopodia during ER depletion, we treated cells with the TRPC1/4/5 agonist, (-)-englerin. It should be noted that TRPC1 had a questionable ability to form channels as a homomer (Xu et al., 2008), (see for review (Dietrich et al., 2014)), but (-)-englerin activated TRPC4 and TRPC5 channels which form heteromeric complexes with TRPC1 (Rubaiy et al., 2018). However, (-)-englerin failed to rescue the filopodia (Figure 2C-D).

The effect of calmodulin inhibitor, W-7, was weaker than that of CALP2 (cf. Figure 2A-B and C-D), so we considered the possibility that CALP2 had multiple targets in addition to calmodulin. CALP2 was designed to bind the EF-hands of calmodulin, but calmodulin is only one of a large family of proteins containing four EF-hands (Zimmer et al., 2013). Although CALP2 could mimic the effect of Ca^2+^ in some experiments (Ten Broeke et al., 2001;Ten Broeke et al., 2004), the current results suggested that CALP2 acted as a calmodulin antagonist since the direction of the effect was replicated by W-7. To determine whether CALP2 and dynasore affected filopodia through similar mechanisms, cells were treated with both agents. These results showed no difference over CALP2 alone (Supplementary Figure 2A), suggesting that the same mechanism was affected.

#### 3.1.4 Vesicle traffic is altered during ER depletion

Finally, another Ca^2+^/calmodulin-activated enzyme, PI3-K, was investigated. Ca^2+^/calmodulin binding to the regulatory p85 subunit of PI3-K is known to release the enzymatically active p110 subunit. Blocking this release with CALP2 would mean the catalytic subunit would be retained in inactive form, which would be duplicated by wortmannin-mediated inhibition of p110. Indeed, its effect was like that of CALP2 (cf. Figures 2A-B and E-F). There are multiple pathways that may be inhibited by wortmannin (see Supplementary Figure 2B), but one would affect endosome traffic, which is regulated by PI3-K classes I and II. By inhibiting class I PI3-Ks, wortmannin may decrease production of phosphatidylinositol 3,4-bisphosphate (PI(3,4)P2), which participates in endosome maturation. The possibility that both CALP2 and dynasore affected the balance of endocytic and exocytotic activity was further explored. One way such an alteration could inhibit filopodia during ER depletion was by inhibiting constitutive exocytosis, which would reduce the size of the membrane pool available for filopodia formation. To test this, cells were treated with brefeldin A. This had no effect on filopodia (Figure 2 G-H), so filopodia disappearance could not be attributed to a shortfall in the pool of membrane available for constitutive exocytosis.

The above results suggested that both dynasore and PI3-K could have affected filopodia by inhibiting endocytosis. Further experiments were done in the framework of a classical two-compartment model of fluid-phase kinetics (Swanson et al., 1985;Heckman et al., 2001). Markers for endocytosed contents enter an exchangeable compartment, from which they are either recycled to the extracellular environment or trafficked into a nonexchangeable compartment. The activation of PKCs by phorbol esters increased the rate of fluid-phase endocytosis, resulting in unbalanced endo- and exocytosis. This trapped considerable amounts of membrane in endocytic compartments (Heckman et al., 1996), which might have explained how elevated [Ca^2+^]_i_ decreased the filopodia. Comparing the HRP uptake of cells in culture medium with that during ER depletion, we found that there was no immediate change in uptake. However, the data represented the amount of uptake of the whole culture, composed mainly of groups and colonies. As shown in Figure 1C, groups of cells lagged behind single cells in showing elevated [Ca^2+^]_i_ in response to ER depletion. Maintaining the cultures for 12 minutes beyond the usual 30-minute ER depletion time decreased the capacity of the exchangeable compartment (Figure 3A). As only single cells were measured when analyzing filopodia prevalence, these data were representative of [Ca^2+^]_i_ in relation to changes in filopodia. In a previous study, the rate of instantaneous marker trafficked into the nonexchangeable compartment was found to be small compared to the amount in the exchangeable compartment. In that case, fluid taken up into the exchangeable compartment was recycled at the rate of ∼228 femptoliters/10^5^ cells/minute. Since efflux from the nonexchangeable compartment was very small, and possibly nonexistent (Heckman et al., 2001), the HRP remaining at the 42-minute time point may have approximated the nonexchangeable content (see below). Cells treated with dynasore in Ca^2+^-free HBBS with CPA showed slightly less marker than those treated for 30 minutes by ER depletion alone. This may have reflected a slower rate of filling the exchangeable compartment in these cells.

**Figure 3.**
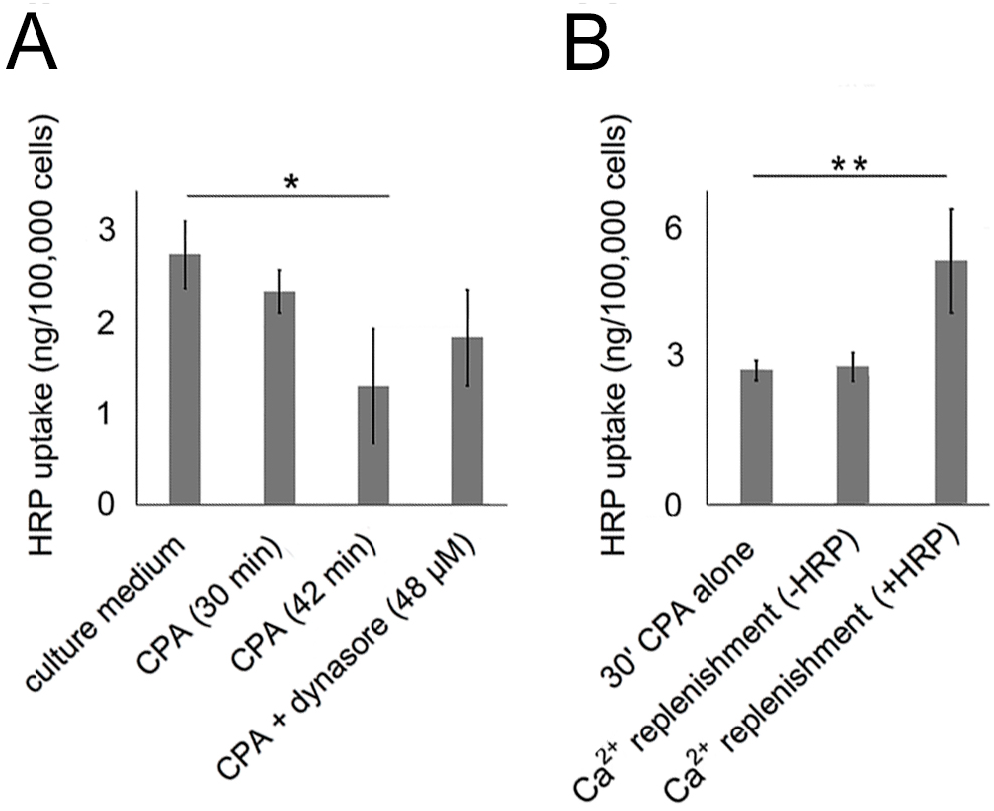
Fluid-phase marker accumulation during ER depletion and Ca^2+^ replenishment. The internalized fluid-phase contents of cultured cells were determined using 1 mg/ml horseradish peroxidase (HRP). Cells were maintained under the conditions designated, then rinsed and lysed in 0.2 % Triton X-100. Results shown are representative of 8 experiments. (A) Cells with marker added for 42 minutes in Ca^2+^-free HBBS with CPA or for 30 minutes in culture medium or Ca^2+^-free HBBS with 5 *μ*M CPA with or without dynasore. Significance by ANOVA, P=0.045, *Treatments differ at P=0.034. (B) Cells maintained in Ca^2+^-free HBBS with 5 *μ*M CPA and 1 mg/ml HRP followed by CPA washout and Ca^2+^ replenishment. Ca^2+^ replenishment was done in the presence or absence of HRP (see procedure in Materials and Methods). Significance by ANOVA, P=0.0054. **P=0.014

The above data indicated that the capacity of the exchangeable compartment shrank during the ER depletion phase. Exocytosis from the RRP may have continued throughout ER depletion, because the HRP content appeared to decrease between 30 and 42 minutes (Figure 3A, columns 2 and 3). To determine the effect of Ca^2+^ replenishment, cells were allowed to internalize HRP during ER depletion, and samples were compared after Ca^2+^ was added in the presence or absence of HRP. As there was no further marker loss on account of Ca^2+^ readdition (Figure 3B), the shrinkage did not continue during this phase. Ca^2+^ replenishment caused an increase in uptake (Figure 3B), indicating the restoration of the exchangeable compartment, however. A large pool of non-secretory exocytotic vesicles is present in the cell, that can serve as a reserve pool of membrane for increasing the capacity of this compartment (see for review, (D’Alessandro and Meldolesi, 2019)).

Initially, the HRP uptake during ER depletion appeared consistent with the hypothesis that membrane tension is a major factor regulating filopodia dynamics (see **1 Introduction**). During ER depletion, the steady-state level of tension may be maintained by endocytosis and limit filopodia formation. This could account for the inverse relationship between endocytosis and filopodia, which was supported by the effects of dynasore and hypertonic sucrose as well as of 1-butanol, a third inhibitor of clathrin-mediated endocytosis which also rescued filopodia (data not shown). On the other hand, the results showing increased marker uptake during Ca^2+^ replenishment in the Ca^2+^ depletion-readdition protocol (Figure 3B), were inconsistent. The filopodia were formed during active endocytosis, but these data are not necessarily irreconcilable with the tension hypothesis. HRP accumulation may have occurred after regeneration of the exchangeable compartment from a reserve pool, followed by rapid reconstitution of the recycling compartment (see **4.1 Relationship of [Ca**^**2+**^]_**i**_ **and Ca**^**2+**^ **fluxes to membrane tension**).

#### 3.1.5 Location of SOCE mediators and aquaporin before and during ER depletion

Previous work in the field showed that STIM activates Orai channels and binds to TRPC1/4/5 channels (see for review (Salido et al., 2011)) which reside in internal vesicles. When we localized STIM1 and TRPC1 with other channel proteins pairwise by immunofluorescence procedures, neither STIM1-TRPC1 nor STIM1-Orai colocalization was common in untreated cells (Figure 4A, STIM1-Orai and STIM1-TRPC1). In contrast, STIM1 and AQP4 were commonly colocalized, and AQP4 was present in both small- and large-sized vesicles with the smaller closer to the cell edge (Figure 4A, STIM1-AQP4). With the exception of AQP4, TRPC1 colocalization with other constituents of vesicles was also rare, including the channel subunit of the VACC, CaV1.2, or caveolin (Figure 4B). TRPC5 was known to reside in vesicles that exchange rapidly with the plasma membrane (Bezzerides et al., 2004), so we determined the extent of TRPC1 colocalization with a marker for the RRP, Vamp2. Their colocalization was less frequent than TRPC1-AQP4 (Figure 4B, TRPC1-AQP4 and TRPC1-Vamp2, Table 1).

**Figure 4.**
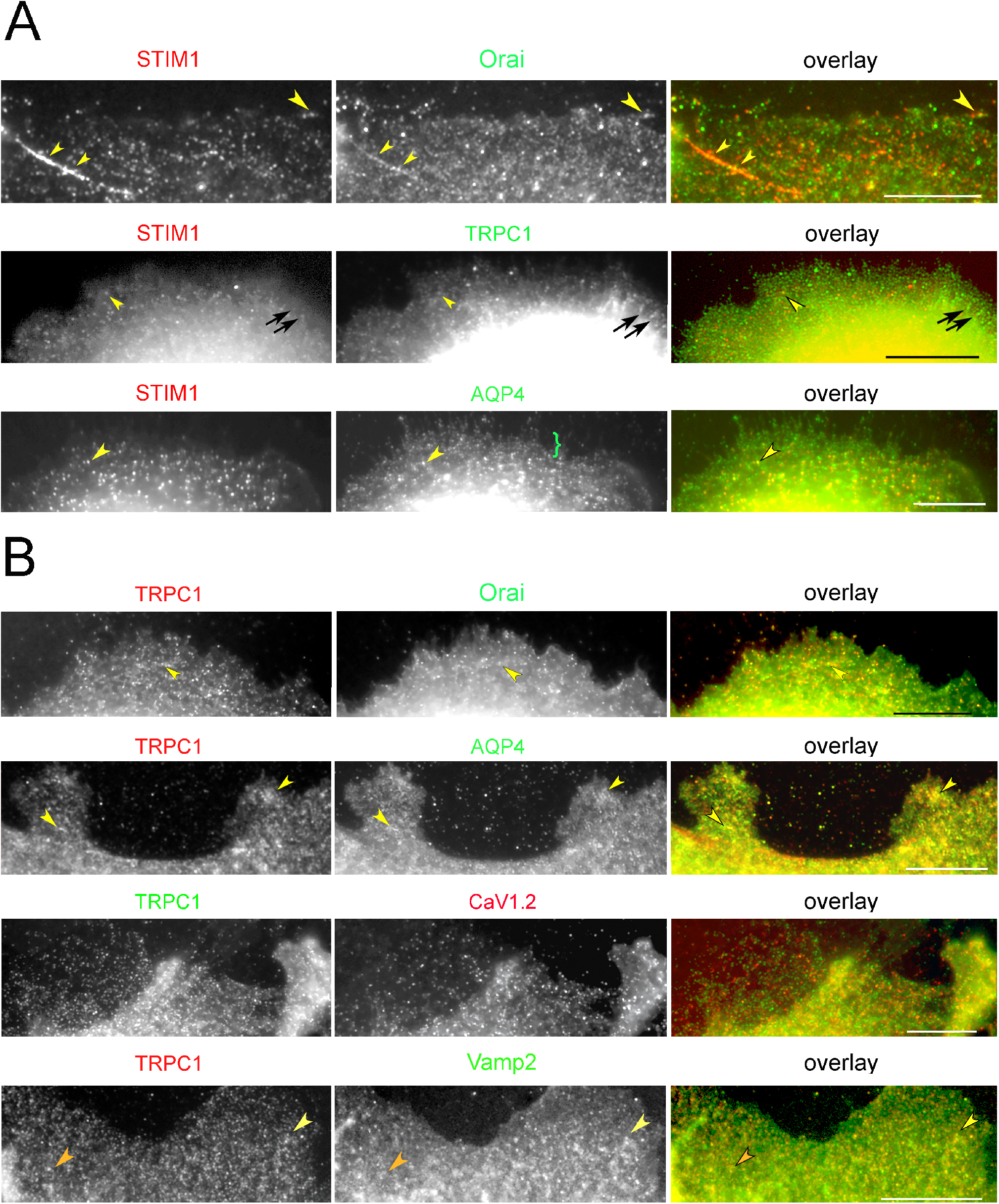
Colocalization of STIM1 and TRPC1 with other proteins in untreated cells. Loci with colocalized proteins are indicated by yellow arrowheads. Coincidence of the labels in higher parts of the cell was ignored because of the addition of fluorescent signals originating from deeper levels in the cell. (A) STIM1 shows little tendency to colocalize with Orai1 or TRPC1 but consistent colocalization with AQP4. TRPC1 is found in linear arrays (black arrows). AQP4 appears in loci of two distinct size classes, with the smaller confined to the cell edge (green bracket). (B) TRPC1-containing loci colocalize with Orai1 and AQP4 but infrequently with the VACC channel subunit, CaV1.2. Areas where the red and green images coincide are mainly in thick portions of the cell where loci are superimposed due to overlapping layers of structure. TRPC1 coincides with Vamp2 in some sites (yellow arrowheads), but in others, diffuse Vamp2 is localized around TRPC1 sites (orange arrowheads). Bars = 10 µm

**Table 1.** Correlation coefficients for proteins paired with STIM1 and TRPC1 in untreated cells*

| Protein coinciding with STIM1 |  | Orai | TRPC1 | AQP4 |
| --- | --- | --- | --- | --- |
| Correlation coefficient |  | 0.509<sup>†</sup> | 0.465<sup>‡</sup> | 0.816<sup>†,‡</sup> |
| Standard error |  | 0.043 | 0.033 | 0.012 |
| Number of images (N) |  | 11 | 17 | 8 |
| Protein coinciding with TRPC1 | CaV1.2 | Orai | Vamp2 | AQP4 |
| Correlation coefficient | 0.552<sup>§</sup> | 0.520 | 0.546<sup>¶</sup> | 0.728<sup>§,\|\|,¶</sup> |
| Standard error | 0.029 | 0.039 | 0.030 | 0.039 |
| Number of images (N) | 10 | 7 | 8 | 8 |

\*Peripheral cytoplasm from images processed by Colocalization Finder (see Materials and Methods).
Significance by one-way ANOVA of comparisons with STIM1 is $P < 0.00001$ . ANOVA with TRPC1 is $P = 0.00031$ .<sup>†</sup>differ by $P = 0.00000$ <sup>‡</sup>differ by $P = 0.00000$ <sup>§</sup>differ by $P = 0.00303$ <sup>||</sup>differ by $P = 0.00046$ <sup>¶</sup>differ by $P = 0.00215$

These observations were confirmed by determining the Pearson correlations for pairs of proteins in the peripheral cytoplasm. Areas similar to those shown in Figure 4 were analyzed with the results shown in Table 1. Table 2 shows the coefficients of the proteins whose correlation with STIM1 or TRPC1 changed significantly during ER depletion or Ca^2+^ replenishment. For example, the correlation for TRPC1-Vamp2 was low in untreated cells as mentioned above, but increased during ER depletion (Table 2). Some proteins showed coefficients with TRPC1 that did not change, including caveolin, Orai, or VACC channel alpha-1 subunit. The Pearson coefficients for these pairs are shown in Supplementary Table S1.

**Table 2.**
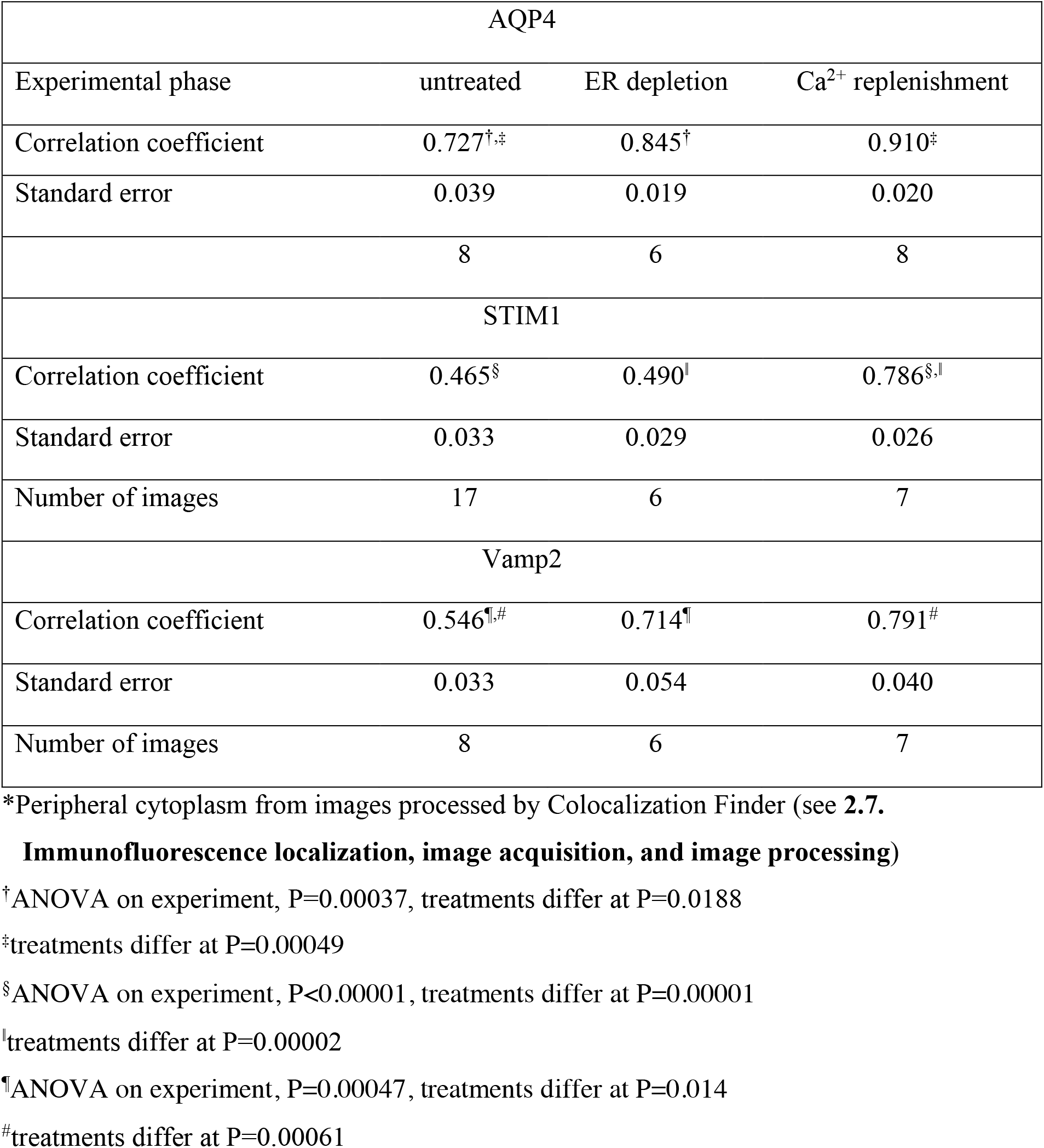
Correlation coefficients for proteins colocalized with TRPC1 before and during SOCE*

While TRPC1 and STIM1 rarely occupied the same compartment in the peripheral cytoplasm, they were found in parallel structures (Figure 5A). TRPC1 was present on filopodia but occupied alternating sites with STIM1 rather than coinciding with it (Figure 5B-C). Colocalization of TRPC1 with β-tubulin demonstrated that TRPC1 was in close proximity to microtubules (Figure 5D). STIM is trafficked as a microtubule end-tracking protein (Grigoriev et al., 2008), and the localizations of TRPC1 suggested that it was also attached but in separate vesicles. In contrast, Orai loci did not form long, linear arrays. Microtubules often terminated near the cell edge but did not extend into filopodia (Figure 5E). In thin portions of cells, the TRPC1-containing loci could be found free of microtubules (Figure 5E and F). The results suggested that TRPC1 and other proteins mainly occupied separate compartments, with the exception of AQP4, which was in separate classes of either TRPC1- or STIM1-bearing vesicles (Table 1).

**Figure 5.**
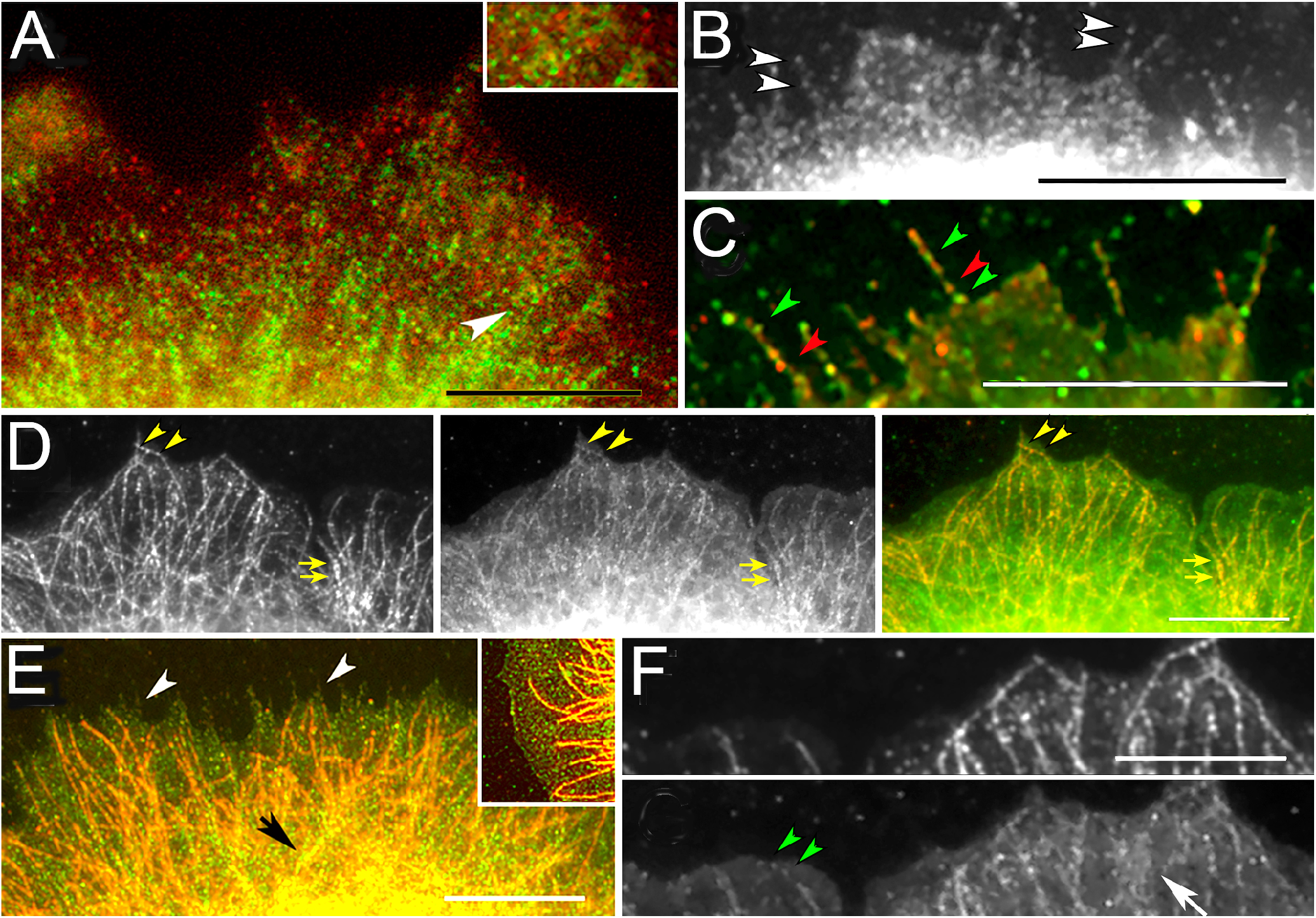
TRPC1 on microtubules and filopodia. (A) Linear arrays of TRPC1 (green) are aligned parallel to features containing STIM1 (red) in an untreated cell. Inset: enlargement of area designated by arrowhead, showing features side by side. (B) TRPC1 loci aligned on filopodia (white arrowheads). (C) TRPC1 loci (green arrowheads) alternating with STIM1 (red) during Ca^2+^ replenishment. (D) Colocalization of β-tubulin (left panel) and TRPC1 (middle panel) in an untreated cell. TRPC1 colocalizes with microtubules bordering the edge (arrowheads) and in the interior (arrows). Panel at right is the overlay. (E) Microtubules (red) labelled by antibody against β-tubulin penetrate to the cell edge but do not enter filopodia (white arrowheads) in an untreated cell. The colocalization of TRPC1 and β-tubulin is apparent in higher parts of the cell (black arrow). Inset: TRPC1 loci (green) free of microtubules (red) at the cell edge. (F) Colocalization of β-tubulin (upper frame) and TRPC1 (lower frame) shows that TRPC1 spots (green arrowheads) extend beyond the ends of microtubules. Interior to the edge, TRPC1 is found in diffuse areas around the microtubules (white arrow). Bars = 10 µm

ER depletion caused very little difference from the patterns shown in Figures 4 and 5, except that STIM1 and Orai were occasionally observed together in plaques. The locations of TRPC1, Orai, and aquaporin channels did not suggest migration to the cell surface during the ER depletion interval. The relationship of STIM1 punctae to Orai is well known and nearly universal (see for review, (Carrasco and Meyer, 2011;Prakriya and Lewis, 2015;Derler et al., 2016)), so our observations are presented in Supplementary Material (Supplementary Figure 3A-B).

### 3.2 Role of SOCE mediators and channels in filopodia formation during Ca^2+^ replenishment

The restoration of extracellular Ca^2+^ causes STIM punctae to detach from sites on the plasma membrane and return to the ER (see for review (Prakriya and Lewis, 2015)). To determine whether this process accompanies the increase in filopodia, STIM1 and the proteins implicated in SOCE were localized again. There were still a few cells showing plaques containing both Orai and STIM1 (Supplementary Figures S3A-B), and the STIM1-Orai correlation in thin portions of the cells did not change (Supplementary Table S1). TRPC1 and STIM1 became colocalized in diffuse areas of the cell surface during Ca^2+^ replenishment, however. These differed in appearance from the STIM1-Orai plaques (cf. Supplementary Figures S3A-B and C). By colocalizing Vamp2 and TRPC1 in cells after Ca^2+^ replenishment, we found a similar pattern. These proteins appeared to be secreted as exosomes in addition to being disseminated on the surface (Supplementary Figures S3D-E). The increased Vamp2-TRPC1 colocalization was confirmed by a significant increase in correlation coefficients over untreated cells (Table 2). Single vesicles containing TRPC1 and Vamp2 were also distributed outside the cell boundary, suggesting that exosomes were released during Ca^2+^ replenishment (Supplementary Figures S3D-E). Untreated, control cells showed little shedding of exosomes (Supplementary Figure 3F). When TRPC1 was colocalized with the voltage-activated calcium channel, CaV1.2, or caveolin, there was no change in their distributions or correlations during ER depletion or Ca^2+^ replenishment. This suggested these proteins were unrelated to filopodia formation, and so the results are shown in Supplementary Materials (Table S1).

Aquaporins participate in polarity determination and directional motility (Karlsson et al., 2011), (see for review (Schwab et al., 2007;Papadopoulos et al., 2008)). Because they conduct water into the space underneath the plasma membrane, which increases the rate of actin polymerization (Karlsson et al., 2013), (see for review (Loitto et al., 2009)), they may facilitate filopodia extension during Ca^2+^ replenishment. When the correlation coefficients between AQP4 and TRPC1 in peripheral areas were determined, their values increased significantly during ER depletion and remained high during Ca^2+^ replenishment (Table 2). To better understand the relationship of SOCE to polarity, we determined which areas of the cells showed a lowered refractive index. Areas near the cell edge, including ruffles and both rounded and pointed protrusions were lower, suggesting that aquaporin was localized at the plasma membrane (Figure 6A). Whereas a specific role for aquaporins in protrusion formation had been demonstrated previously (Hara-Chikuma and Verkman, 2006;Karlsson et al., 2013), these data suggested they had a general effect on the edge of 1000 W cells (Figure 6A).

**Figure 6.**
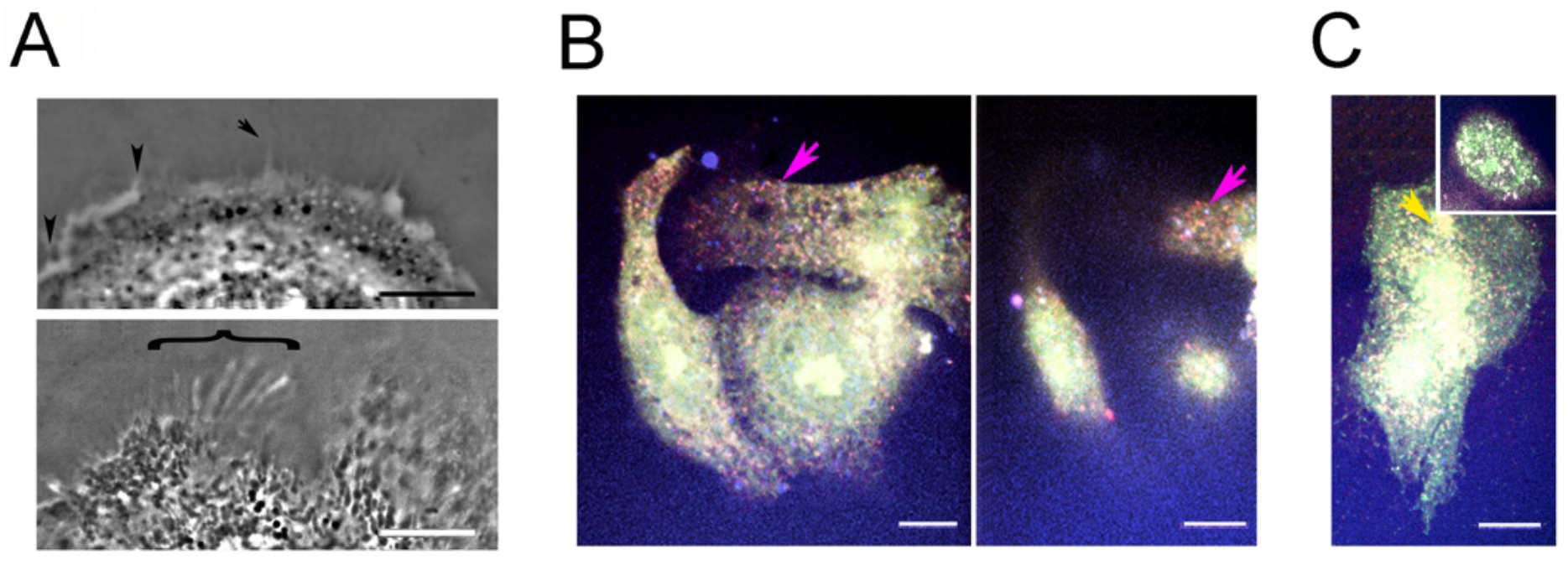
Localization of aquaporin and STIM1-labelled loci in relation to channels. (A) Areas with low refractive index as indicated by phase density, representative of 3 experiments. (Top) Low density areas are found immediately under the plasma membrane (arrowhead), in vesicles near the cell edge, in elevated portions of the cell, and in pointed protrusions (arrow). (Bottom) Linear structures with low density are present in the lamellipodium (bracket). (B, C) Confocal planes from cells after Ca^2+^ replenishment and colocalization with antibodies against STIM1 (blue), Orai (green), and TRPC1 (red). (B, Left) Ventral plane with diffuse Orai (green) and numerous areas of STIM-TRPC colocalization in magenta (arrow). (Right) Dorsal plane of the same cells, (C) Ventral plane of another area showing Orai colocalized with TRPC1 (yellow arrowhead) and areas of STIM1-TRPC1 colocalization (magenta foci). Inset: Nucleus of the same cell showing the diffuse distribution of Orai (green) and colocalization of all three proteins (white). Bars =10 µm

To determine whether all three proteins mediating SOCE coincided, they were colocalized using antibodies against STIM1 (blue), Orai (green), and TRPC1 (red). These results confirmed that STIM1-TRPC1 colocalization was common (Figure 6B, magenta arrows), while STIM1 and Orai coincided much less frequently (cf. Figure 4B and cyan in Figure 6 and Supplementary Figure 3I). The frequency of TRPC1-STIM1 colocalization may have occurred because their content was much higher than endogenous Orai. TRPC1-Orai colocalization occurred in more central or elevated parts of the cell (cf. Figure 6C and Supplementary Figure S3G-I (yellow arrowheads), as did the colocalization of all three proteins (Figure 6C, white areas).

#### 3.2.1. SOCE alters size distributions of vesicles bearing STIM1 and channel proteins

The evidence above indicated that TRPC1 was in vesicles attached to microtubules. Because the localization of TPRC5 to vesicles of the RRP is well-known (Bezzerides et al., 2004), and TRPC1 resides in internal vesicles, and it is brought to the plasma membrane by Rab4-dependent recycling (Cheng et al., 2011;de Souza et al., 2015), it was surprising that TRPC1 and Vamp2 were not completely colocalized (Table 2). To gain further insight into TRPC1 vesicle movements, we analyzed the radii of loci in images obtained by localizing the vesicle-bound proteins. This was done by segmenting the discrete loci and evaluating the ones that had a circular form (see **2.11 Analysis of particles’ dimensions after staining by indirect immunocytochemistry**). The resulting distributions showed peaks in different ranges as shown in Figure 7A-E.

**Figure 7.**
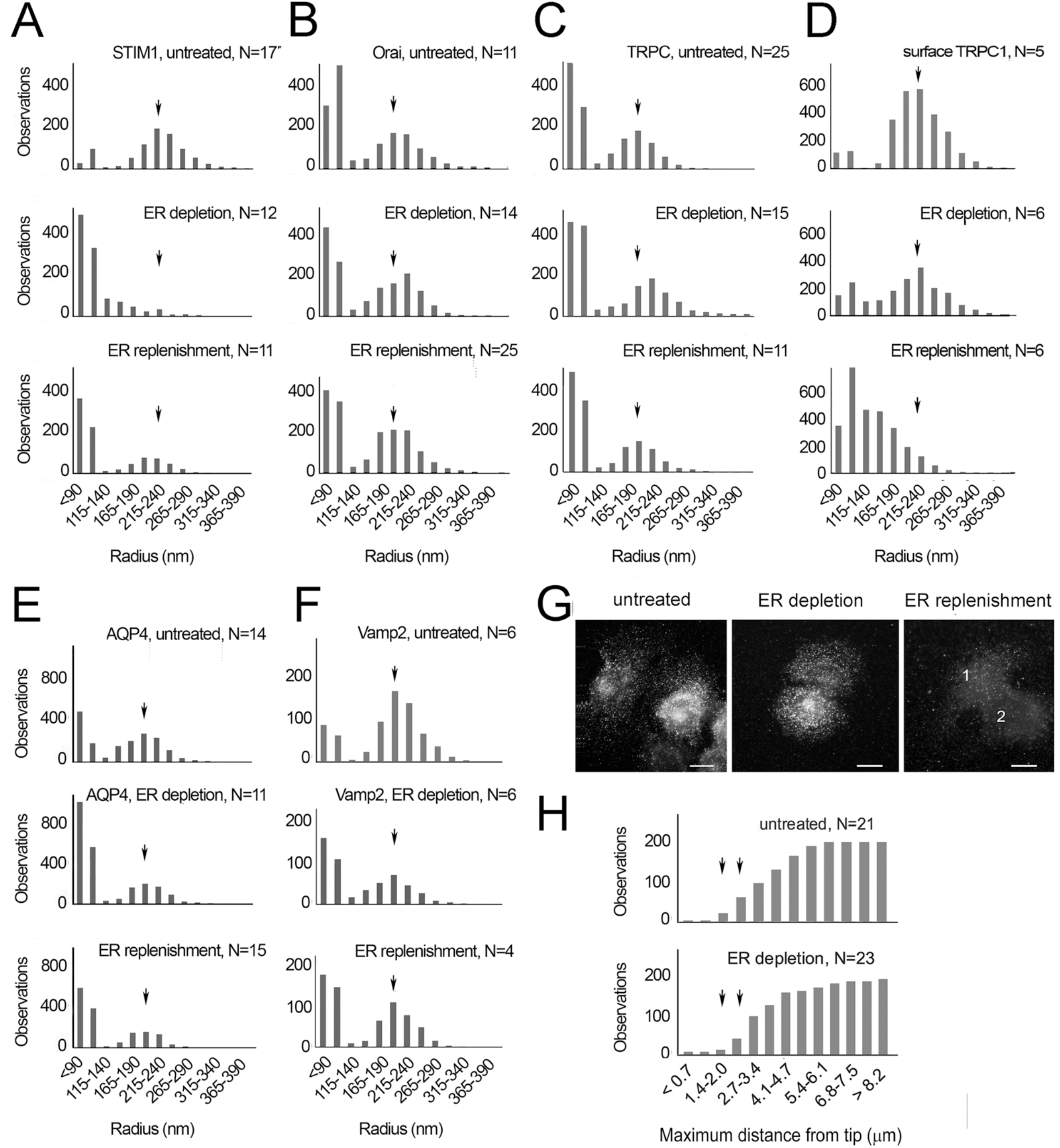
Characteristics of vesicles displaying intrinsic membrane proteins, STIM1, Orai, TRPC1, AQP4, and Vamp2. Results represent 3-8 experiments. (A-F) The peak radius for each vesicle size distribution in the untreated cell sample is indicated by an arrow and the same mark applied to samples collected during ER depletion and Ca^2+^ replenishment. (A) STIM1 peak at 240 nm, (B) Orai peak at 210 nm, (C) TRPC1 peak at 200 nm, (D) Cell surface TRPC1 peak at 215 nm, (E) AQP4 peak at 205 nm. (F) Vamp2 peak at 205 nm, (G) Cells were fixed but not permeabilized either after no treatment or after the Ca^2+^ depletion part of the protocol, and the samples processed for TRPC1 localization using the antibody against the domain exposed on the cell surface. TRPC1 is diffuse on the surfaces of cells 1 and 2. (H) Cumulative distribution of TRPC1 loci in untreated cells compared to cells during ER depletion. The circle within which loci are found near each filopodium is represented as its diameter. Loci within 2.7 µm of the filopodia tip are slightly fewer during ER depletion. Statistics on the distances are untreated (mean=3.27, S.E.M. 0.091 µm, 196 loci) and ER depletion (mean=3.78, S.E.M. 0.13 µm, 187 loci). N = number of images analyzed

During ER depletion, STIM-bearing portions of the ER are found at the plasma membrane. This was confirmed by the appearance of STIM1-Orai plaques (Supplementary Figure 3A-B) and was also reflected in the smaller diameters of STIM loci (Figure 7A). Numerous loci of 140 nm or less were found (Figure 7A), representing the well-known punctae in the plasma membrane described elsewhere (Grigoriev et al., 2008;Carrasco and Meyer, 2011;Asanova et al., 2014;Liu et al., 2015). The radii of Orai-and TRPC1-bearing vesicles changed in the opposite direction, increasing slightly. For example, the TRPC1 peak increased from ∼200 to ∼220 nm. These distributions reverted to their original peak values after Ca^2+^ replenishment (Figure 7A-C). Despite these changes, TRPC1-Orai colocalization remained uncommon (Figure 4B, TRPC1-Orai and Supplementary Figure 3G-I), and their Pearson correlations were unchanged throughout the Ca^2+^ depletion-readdition protocol (Supplementary Table S1). The shrinkage of the exchangeable compartment was accompanied by slight increases in the TRPC1 and Orai peak areas. Both recovered after Ca^2+^ restoration (cf. Figure 3B and 7B-C). The data suggested that the original TRPC1 distribution represented vesicles of the exchangeable compartment, and their apparent sizes increased when the membranes merged with a reserve pool of larger-sized vesicles.

The TRPC1 loci of Figure 7C represented a mixture of intracellular vesicles and surface sites. When we imaged the latter sites, using an antibody directed against the extracellular portion of the TRPC1 channel (**2.7 Immunofluorescence localization, image acquisition, and image processing)**, their peak diameter was slightly larger than the mixed intracellular and surface loci. The 215 nm peak remained the same during ER depletion (Figure 7D) but was dramatically reduced during Ca^2+^ replenishment. Aquaporins were also present on the plasma membrane (see **3.2 Role of SOCE mediators and channels in filopodia formation during Ca**^**2+**^ **replenishment**). AQP4 was colocalized with TRPC1 and STIM1 during ER depletion and Ca^2+^ replenishment (Tables 1 and 2). The AQP4 size distribution was similar to that of TRPC1 (cf. Figure 7C and E), because the sites analyzed were near the cell edge. A slight reduction during ER depletion, resembling that of STIM1-bearing vesicles (cf. Figure 7A and E), suggested that they may be translocated with STIM. For Vamp2, a slight increase in small (<140 nm) particles suggested it may be secreted during ER depletion and Ca^2+^ replenishment (cf. Figure 7E and 7F). This was accompanied by a significant increase in the correlation coefficient of Vamp2 with TRPC1 (Table 2).

TRPC1 appeared on the cell surface before and during ER depletion as mentioned above, but dissemination was also apparent after Ca^2+^ replenishment when localizations were done with the rabbit anti-TRPC antibody which localized both intracellular and extracellular sites (cf. Figure 7G and Supplementary Figure 3C). At the same time, the peak size distribution of STIM1 loci reverted to a pattern resembling that of untreated cells (Figure 7A). The results suggested that STIM1 loci were dispersed in plaques during ER depletion, followed by TRPC dispersion after Ca^2+^ readdtion. To test whether STIM and TRPC1 were translocated together, we sought further evidence that TRPC surface exposure was unchanged during ER depletion. We measured the number of TRPC1 clusters localized near each filopodium, using the antibody against the extracellular portion of the molecule to determine whether the density of sites increased. There was a slightly lower TRPC1 representation in areas surrounding the filopodia, indicating little change in surface TRPC1 (Figure 7H). Although cells undergoing Ca^2+^ replenishment were also analyzed, this was uninformative because the observations extended over longer distances due to filopodia extension (data not shown). Because TRPC1 dissemination was associated with filopodia formation, we also explored the possibility that dynasore or CALP2 treatment had caused TRPC dissemination during ER depletion. Comparing the CALP2- or dynasore-treated with cells undergoing ER depletion alone, we found greater representation of external TRPC1 loci with a radius <140 nm (Supplementary Figure 4A-D). The results were unlikely to be due to increased exocytosis of TRPC1, as neither Ca^2+^/calmodulin nor dynamin directly affected exocytosis. Thus, TRPC1 was thought to be trapped in the plasma membrane because the treatments inhibited internalization in endosomes. Because internal TRPC1 is largely in the RRP, the data suggest that its surface concentration is regulated by rapid exchange with the RRP. Therefore, we proposed that this limitation on the TPRC1 concentration is partially relieved by CALP2 or dynasore.

#### 3.2.2 Filopodia formed during Ca^2+^ replenishment depend on TRPC1/4/5

As the above data suggested that TRPC1 vesicles as well as surface clusters were disseminated during Ca^2+^ restoration, we postulated that surface TRP molecules were important for filopodia extension. A number of channel inhibitors were tested to determine whether any blocked filopodia formation. SKF96365, as well as a specific antagonist of TRPC1, 4, and 5 heteromeric complexes, pico145 (Rubaiy et al., 2017), inhibited filopodia. Nifedipine, a VACC inhibitor, was slightly inhibitory (Figure 8A-B). The results showed that TRPC1, 4, and/or 5 were important for filopodia formation. To determine whether the other major subfamily of TRPC1 isoforms, TRPC3/6/7, contributed, Ca^2+^ was restored in the presence or absence of dioctylglycerol (DOG), which was expected to activate them. This had no impact on the filopodia (Figure 8C-D), and thus, no role for the TRPC3/6/7 subfamily in filopodia could be confirmed.

**Figure 8.**
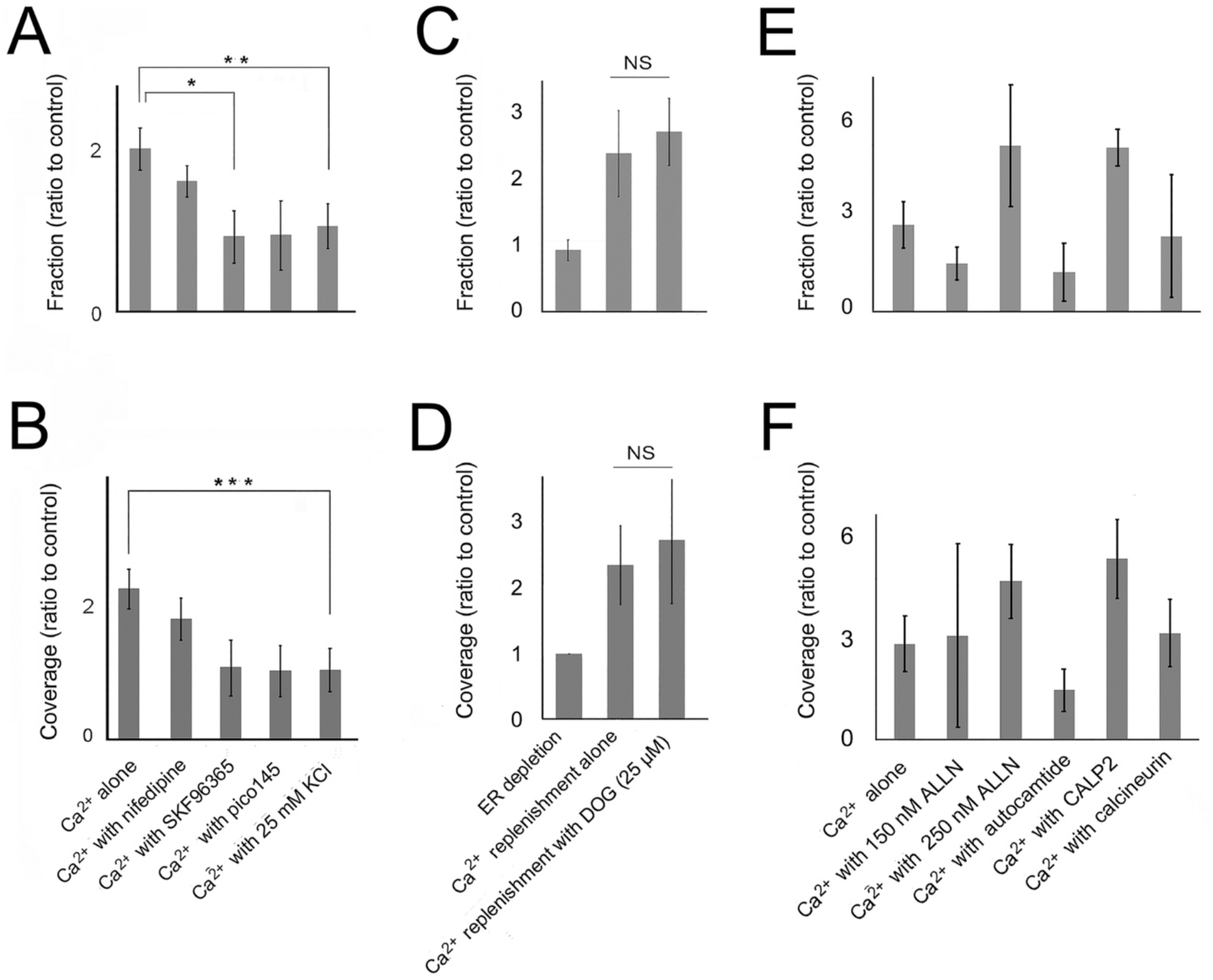
Ca^2+^ channels and effectors implicated in net filopodia formation. (A-F) Cells in Ca^2+^-free HBSS were treated with CPA, rinsed, and then treated further by replenishment of the extracellular Ca^2+^ prior to fixation. Results are representative of 3-9 experiments. (A-B) Filopodia prevalence after Ca^2+^ replenishment alone (Ca^2+^ alone) or with 8 µM nifedipine, 10 µM SKF96365, 15 nM pico145, or 25 mM KCl. (A) Significance by ANOVA, P=0.0075, *P=0.032, **P=0.047. (B) Significance by ANOVA, P=0.012, ***P=0.027. (C, D) Filopodia prevalence of cells treated with replenishment alone (Ca^2+^ alone) or with 25 µM dioctylglycerol (DOG), (E, F) Filopodia prevalence of cells treated with replenishment alone (Ca^2+^ alone), ALLN, 10 µM autocamtide-2 inhibitory peptide, 20 µM CALP2, or 20 µM calcineurin inhibitory peptide. Significance by ANOVA, P=0.165.

#### 3.2.3 Exocytosis and stimulus-coupled secretion during extracellular Ca^2+^ replenishment

As the evidence suggested that filopodia extension was accompanied by TRPC1 secretion along with part of the cell’s Vamp2 (Figure 7G, Supplementary Figure 3C, Table 2), we reexamined the effect of the inhibitors that were used during ER depletion (Figure 2A-D). The effects of ALLN and CALP2 were still positive but fell short of being statistically significant (cf. Figure 2A-B and 8E-F). Their effects were opposite in direction from those of hypertonic sucrose and dynasore (cf. Figure 8E-F and 9A-B). Although we proposed above that blocking endocytosis during ER depletion caused TRPC1 to be trapped (see **3.2.1. SOCE alters size distributions of vesicles bearing STIM1 and channel proteins**), a similar trapping effect during Ca^2+^ replenishment became inhibitory (cf. Figures 2C-D and 9A-B). The simple explanation for this reversal was that Ca^2+^ replenishment enhanced the rate of vesicle exchange with the RRP. When recycling was inhibited by dynasore, the RRP rapidly diminished, because it could not be replenished from the plasma membrane. This suggested that filopodia formation during Ca^2+^ replenishment was sustained by vesicle recycling. Moreover, the addition of membrane alone from exocytosis could not maintain extension, suggesting that surface tension was insufficient to enable filopodia extension in the dynasore-treated cells.

Translocation of TRPL, a canonical TRP channel of Drosophila, to the plasma membrane had required a sustained influx of Ca^2+^ (Richter et al., 2011). Results from other laboratories also suggested a role for stimulus-coupled secretion, which relies on Ca^2+^ influx through the VACCs (see for review (Mears, 2004;Seino and Shibasaki, 2005)). In the commonly used model of transmitter release from presynaptic neurons, rapid exocytosis relies on raising [Ca^2+^]_i_ to micromolar levels in domains surrounding the voltage-gated Ca^2+^ channels. Previous evidence, showing that VACCs existed in airway epithelial cells in situ (Ten Broeke et al., 2001), was consistent with our data showing that 25 mM KCl in the Ca^2+^-replete medium increased [Ca^2+^]_i_ (Figure 1F). To determine the role of this secretory mechanism, we depolarized the cells with KCl during Ca^2+^ restoration. This was expected to generate areas of high, local Ca^2+^ influx in the vicinity of the VACCs, but it inhibited filopodia (Figure 8A-B). In KCl-treated cells, TRPC1-bearing vesicles appeared to be larger and more centrally located when imaged after immunolocalization (data not shown).

With regard to filopodia, Ca^2+^ influx through VACCs had a profoundly different effect from Ca^2+^ influx through Orai and/or TRPC channels. In chromaffin cells, the rates of exocytosis induced by depolarizing pulses and VACC currents were much higher than those dependent on SOCE (see for review (Putney and Parekh, 2005)). We used a classical inhibitor of VACCs, nifedipine, to determine whether it would rescue filopodia. They were slightly increased when nifedipine was applied together with KCl, but greatly increased when cells were treated with KCl in Ca^2+^-free media (Figure 9C-D). As large Ca^2+^ influxes were essential for stimulus-coupled secretion, these data showed that the VACC-mediated inhibition of filopodia depended on extracellular Ca^2+^ as well as on exocytosis. If the exchangeable compartment was reconstituted rapidly upon Ca^2+^ replenishment, as proposed above, the size of the replenishing pool could be restricted by blocking constitutive membrane transport from the ER to downstream compartments. Tests with brefeldin A confirmed that it was inhibitory to filopodia formation during Ca^2+^ replenishment (Figure 9E-F). This result suggested that the size of the replenishment pool upstream of the RRP was limited by brefeldin A exposure, leading to inhibition of filopodia extension after Ca^2+^ replenishment.

**Figure 9.**
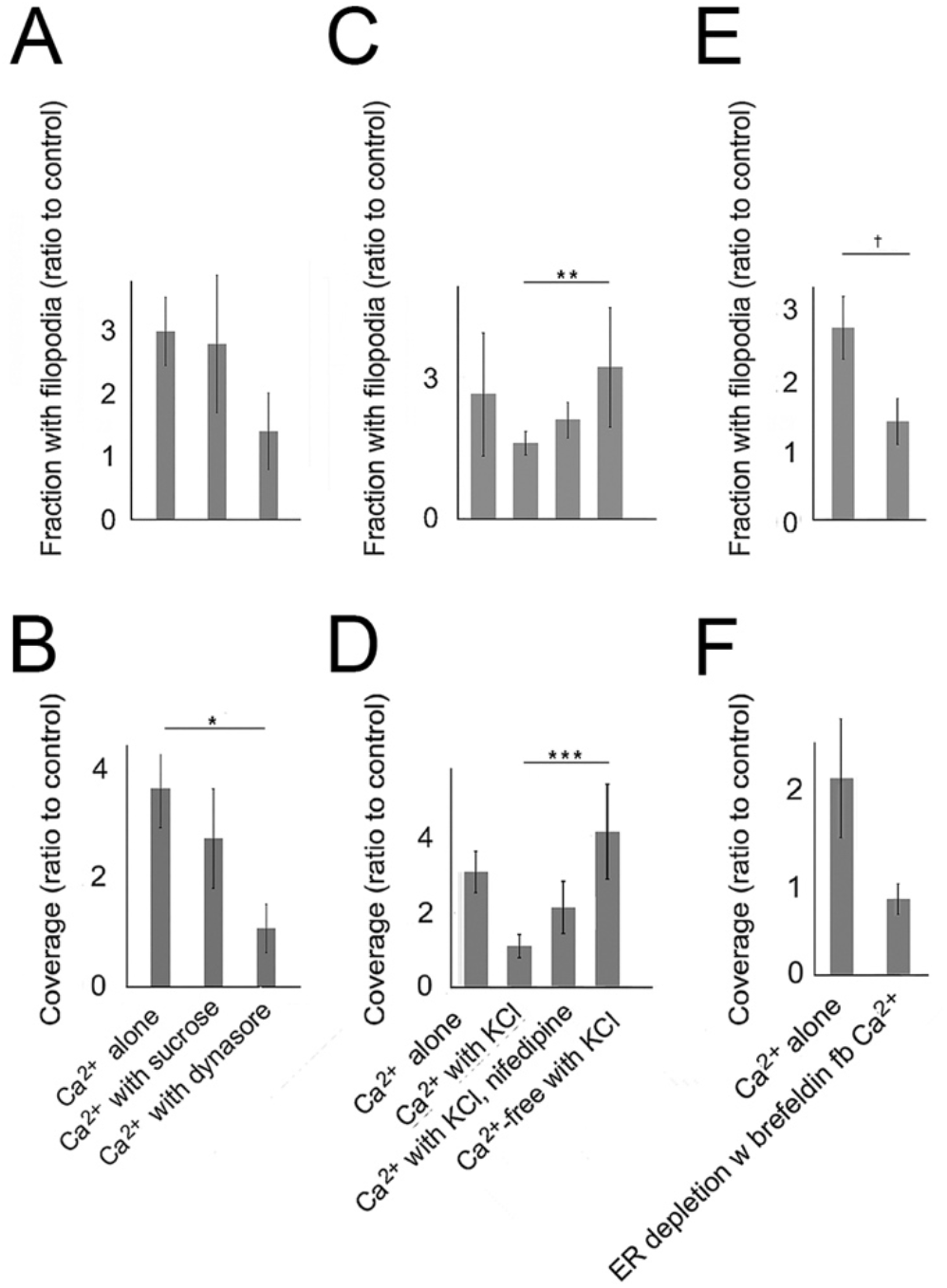
Effects of endocytic and exocytotic inhibitors on filopodia prevalence after Ca^2+^ replenishment. Results shown are representative of 4-9 experiments. (A-D) Filopodia prevalence after Ca^2+^ alone (Ca^2+^ alone) or in the presence of various inhibitors, (A, B) Ca^2+^ replenishment in the presence or absence of 0.4 M sucrose or 48 µM dynasore. (A) Significance by ANOVA, P=0.153, (B) Significance by ANOVA, P=0.0034, *P=0.038. (C, D) Ca^2+^ replenishment in the presence or absence of 25 mM KCl with and without nifedipine or 25 mM KCl in Ca^2+^-free HBSS (Ca^2+^-free) (C) Significance by ANOVA, P=0.049, **P=0.033. (D) Significance by ANOVA, P=0.022, ***P=0.012. (E, F) Ca^2+^ replenishment in samples pretreated during ER depletion with or without brefeldin (20 µM). (E) ^†^Treatments differ at P=0.044.

Altogether, the results suggest that the depletion part of the protocol profoundly decreased the size of the RRP and thereby slowed TRPC1 trafficking. The data of Figure 7G, as well as Supplementary Figure 4C-D suggest that TRPC1 is present on the surface during ER depletion but is not homogeneous as it is during Ca^2+^ replenishment. If Ca^2+^ replenishment reverses the shrinkage of the exchangeable pool and restores membrane recycling, as we propose, the positive effect of these changes on filopodia may depend on dissemination of membrane constituents. Stimulus-coupled secretion may perturb the restoration and thereby interfere with its positive effect on filopodia.

If replenishing the extracellular Ca^2+^ caused an imbalance in exocytosis and endocytosis, favoring the exocytosis of TRPC, we would expect the same inhibitors of endocytosis as used during ER depletion to have a lesser effect on filopodia formation. Indeed, not only did these inhibitors no longer stimulate filopodia formation but their effect was inhibitory (cf. Figures 3C-D and 9F). Although the effects were opposite in direction to that found during ER depletion, they did not rise to a level of statistical significance. If endocytosis in this phase helped replenish the RRP pool, as the data seemed to indicate, recycling of exocytic vesicles through endocytosis may be required to sustain filopodia extension. Since TRPC1 was disseminated on the cell surface after treatment with CALP2 during depletion, and we assume there was little or no effect on exocytosis, a blockade of endocytosis of RRP vesicles was probably responsible for the CALP2-induced filopodia formation.

## 4 Discussion

TRPC channels and the Ca^2+^ sensor, STIM, are known to participate in polarity determination (see **1 Introduction**), but the mechanisms linking them to polarity were poorly understood. We confirmed that Ca^2+^ deprivation in serum-free medium left both [Ca^2+^]_i_ and filopodia unchanged. With CPA treatment for 30 minutes in the absence of Ca^2+^ and signaling constituents, the driving force for SOCE was fully developed, and TRPC1 channels were mobilized to the plasma membrane by Ca^2+^ replenishment. This confirmed reports that TRPC1 channels were mobilized to the plasma membrane from intracellular compartments (see **3.2.1. SOCE alters size distributions of vesicles bearing STIM1 and channel proteins**). It was suggested previously that local Ca^2+^ entry was required for this process, and that TRPC1 was not rapidly reinternalized upon removal of the Ca^2+^-containing medium (Cheng et al., 2011). A large difference in mechanisms was revealed by comparing previous studies with the current experiments, however. Here, replenishment of the extracellular Ca^2+^ acted in the absence of signaling from receptors but in previous studies, TRPC5/6 were mobilized immediately after receptor ligation (Bezzerides et al., 2004;Monet et al., 2012).

Filopodia also increased rapidly after CPA treatment in the current studies, and this occurred without the possibility of Ca^2+^ influx, as the extracellular media were Ca^2+^-free. Cells may have continued processing signals over this short interval, meaning that the conditions may have replicated the immediate responses mentioned above.

The processes of STIM migration to the cell surface, Orai activation, and exocytosis of TRPC-bearing vesicles remained intact despite the absence of extracellular ligands that could occupy receptors. Ca^2+^ replenishment activated both TRPC1 dissemination in the plasma membrane and filopodia extension. Although this suggested a close relationship, any one of the three stages of filopodia dynamics might have been affected, i.e. initiation, extension, or retraction. Initiation typically requires activation of myosin X by the PI3-K product, PI(3,4,5)P3, at sites where actin-related proteins are also assembled (He et al., 2017). During ER depletion, wortmannin, which is a potent PI3-K inhibitor, did not decrease filopodia but rather increased them. This suggested that they were poised for extension throughout the process. Thus, the conditions of these experiments only appeared to affect extension and retraction. The results relevant to these stages are discussed under separate headings below. The net retraction that accompanied ER depletion was consistent with expectations from the tension hypothesis (see **4.1 Relationship of [Ca**^**2+**^]_**i**_ **and Ca**^**2+**^ **fluxes to membrane tension**). Exocytosis from the RRP appeared to be stimulated by Ca^2+^ influx itself (see **4.2 Restoration of extracellular Ca**^**2+**^ **enables exocytosis of TRPC-bearing vesicles**). We propose that Orai functions by directing small Ca^2+^ currents into the space beneath the plasma membrane where TRPC-bearing vesicles are concentrated, and this stimulates TRPC exocytosis (Figure 10).

**Figure 10.**
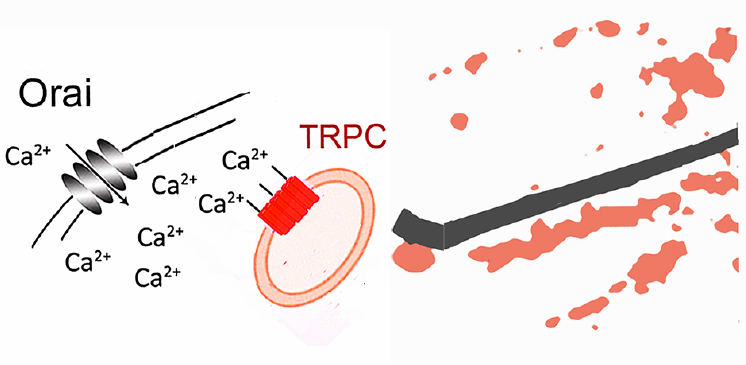
Model for the role of Orai in exocytosis of TRPC-bearing vesicles from the RRP, explaining how filopodia respond to Ca^2+^ in the absence of a Ca^2+^-binding structural protein. The model proposes that the vesicles are retained under the plasma membrane by attachment to a microtubule (black), and their release is stimulated by Ca^2+^. On the contrary, the lack or Orai alignment in linear arrays suggested that it was not mobilized to the cell surface on microtubules.

### 4.1 Relationship of [Ca^2+^]_i_ and Ca^2+^ fluxes to membrane tension

The cell’s ability to make protrusions is limited by surface tension (see 1 Introduction). As endocytic activity was expected to increase the tension on the surface by limiting the area of the plasma membrane, it would oppose the force exerted by the filopodium tip. Our evidence was consistent with this hypothesis, because inhibitors of endocytosis blocked net filopodia retraction during ER depletion. However, direct evidence of endocytic activity from HRP uptake experiments showed no enhancement of endocytosis over the interval of ER depletion. On the contrary, marker uptake declined. The results, taken together, suggested that exocytosis continued while endocytosis was slowed. Exposure to dynasore or hypertonic sucrose slowed endocytosis still further and allowed expansion of the plasma membrane. This added membrane relieved tension on the cell surface and allowed more filopodia to be extended.

Markers like HRP are internalized into a compartment that allows rapid exchange with the extracellular medium. While this exchange continues during ER depletion, the trend for decreasing HRP content suggested that the volume of the compartment was shrinking. Shrinkage was supported by two findings that also suggested retraction of the RRP into the replenishment pool. The radii of the Orai- and TRPC1-bearing vesicles increased slightly (Figure 7), and the distance between the filopodia tip and TRPC1-bearing loci in the cytoplasm also increased (Figure 9H). While the majority of HRP is taken up into an exchangeable compartment, small amounts are trafficked into a nonexchangeable compartment consisting of late endosomes, lysosomes, and trans-Golgi vesicles (Heckman et al., 2001). As the exchangeable contents would undergo efflux when medium without marker was provided after the ER depletion interval, the remaining marker was in the nonexchangeable pool. We propose that the retraction of the RRP compartment led to an increase in surface tension and a failure of filopodia extension. This could be one factor, if not the main factor, accounting for net filopodia retraction. The tension hypothesis is further discussed in section 4.2.

### 4.2 Restoration of extracellular Ca^2+^ enables exocytosis of TRPC-bearing vesicles

During Ca^2+^ replenishment, cells regenerated the exchangeable compartment and renewed their uptake of HRP. In excitable cells, the opening of VACCs allows Ca^2+^ entry, which elevates its levels in nanodomains centered on VACCs (see **3.2.3 Exocytosis and stimulus-coupled secretion during extracellular Ca**^**2+**^ **replenishment**). The resulting wave of exocytosis is followed by compensatory endocytosis. As similar increases in [Ca^2+^]_i_ in the presynaptic terminal following an action potential (Balaji et al., 2008;Yao and Sakaba, 2012), (see for review (Hallermann, 2014;Wu et al., 2014)) precede endocytosis, this could be an alternative model for the renewed HRP uptake (Figure 3). This was a more complex explanation, but it would make the observations consistent with the tension hypothesis. The addition of the newly secreted membrane to the plasma membrane would contribute to the area of the plasma membrane and relieve tension, favoring filopodia formation. When we tested whether endocytosis could have been activated by stimulus-coupled secretion, as it was in the presynaptic terminal, results with KCl contradicted this interpretation. Thus, we concluded that HRP uptake resulted from membrane recycling, and this is closely related to net filopodia extension.

The increase in the size of the RRP led to increased amounts of TRPC1 at the cell surface after Ca^2+^ readdition (Figures 6C, 7D). The current results showed that this type of exocytosis was SOCE-dependent and not VACC-dependent. The hypothesis that there was increased recycling between the exchangeable compartment and the extracellular medium was supported by results with hypertonic sucrose and dynasore. They would block the accelerated recycling upon Ca^2+^ readdition and therefore would inhibit net filopodia extension, consistent with our observations. Moreover, exposure to brefeldin during ER depletion inhibited subsequent filopodia formation upon Ca^2+^ replenishment (Figure 9). Brefeldin A would affect the size of the RRP in a similar way as the inhibitors. In the case of brefeldin, however, the size of the upstream replenishing pool was restricted by limiting constitutive secretion. In this context, it should be noted that the rate of RRP replenishment may depend on Ca^2+^/calmodulin, as it does in the presynaptic nerve terminal (Wu et al., 2009). If the RRP is also responsive to Ca^2+^/calmodulin in epithelial cells, the first phase of SOCE may enlarge the RRP compartment and thereby contribute to TRPC1 exocytosis after Ca^2+^ readdtion.

### 4.3 Polarity, PI3-K/PTEN axis, and Ca^2+^

To fully understand how filopodia responded to Ca^2+^, it was necessary to address how their prevalence was affected by [Ca^2+^]_i_ as well as Ca^2+^ fluxes. Previous work detailing how filopodia responded to spontaneous Ca^2+^ transients in dendrites, had shown that low intracellular Ca^2+^ was required to initiate extension (Lohmann et al., 2005). In many studies of growth cone motility, however, filopodia extension followed an elevation of [Ca^2+^]_i_ (see for review (Ademuyiwa, 2019)). Formation was permitted when [Ca^2+^]_i_ was either low and rising or high and falling – a relationship summarized earlier by Kater and coworkers (Kater et al., 1994) as a “bell-shaped” pattern. In the current studies, both patterns were observed. Filopodia were formed after Ca^2+^ readdition when [Ca^2+^]_i_ was high and falling, but also during ER depletion, when [Ca^2+^]_i_ was low and rising. The discovery that extension required recycling of vesicles from the RRP now provides an explanation. While some RRP vesicles were poised and immediately available for exocytosis as [Ca^2+^]_i_ levels rose, others required a maturation process to ready them for exocytosis. Maturation would be favored under conditions of high [Ca^2+^]_i_. This interpretation supports the hypothesis of Gallop (Gallop, 2020), holding that critical molecules must be exchanged with the plasma membrane in order to ensure filopodia extension. Although tension was sufficient to explain net filopodia retraction during ER depletion, net formation could not easily be explained by the tension hypothesis. Membrane, presumably added to the surface by stimulus-coupled secretion, would relieve surface tension but did not have the same effect as Ca^2+^ readdition.

The issue of how TRP channels are implicated in polarity and chemotaxis is important, because TRPC5 and members of the TRPM and TRPV subfamilies have been implicated in the cancer cell’s ability to migrate and metastasize (see for review (Canales et al., 2019). The proposed relationship among Ca^2+^/calmodulin, RRP capacity, and filopodia, suggested above, may be tested readily with results bearing on the role of polarity in tumor promotion. Despite the fact that front-rear determination is usually established by a gradient of PI3-K and PTEN, (see for review (Mazel, 2017)), and filopodia are concentrated at the leading edge, PI3-K activity did not appear to enhance filopodia in the current experiments. Our results suggested the opposite, that inhibition of PI(3,4,5)P3 production by wortmannin enhanced filopodia. That its effect was mimicked by inhibitors of endocytosis such as dynasore, as shown in Figure 2, suggested that endocytosis was the target during ER depletion when [Ca^2+^]_i_ was low. Its effect also mimicked that of dynasore during Ca^2+^ readdition (data not shown), suggesting that the target now became rapid recycling. While the data overall reinforced the idea that proteins are introduced into the membrane to “seed” filopodia, consistent with Gallop’s hypothesis (Gallop, 2020), they also bear on the larger question of membrane flow in directional locomotion. A proposed scheme in which endocytosis occurs randomly on the cell surface and exocytosis at the front of the cell (Bretscher, 1996) has yet to be confirmed. Our results further complicate this hypothesis by introducing the recycling compartment. Thus, further experimentation is needed to test and verify the conclusions and fully clarify the role of polarity determinants in cell orientation and directional persistence.

## 5 Conclusions

The results suggest that SOCE regulates the size of the RRP in epithelial cells, and vesicle recycling is the immediate mechanism affecting filopodia extension. Dissecting this mechanism further will be essential to understand how calcium ions contribute to cell motility.

## 7 Author Contributions

CAH and OMA conceptualized the studies and developed the methodology. CAH, OMA, and MLC did the investigations. CAH and MLC provided resources. MLC developed and applied software. CAH and MLC validated the studies. The original draft was written by OMA and CAH; the final draft was reviewed and edited by CAH, OMA, and MLC.

## 9 Non-standard abbreviations

AQP4: aquaporin isoform
CALP2: calcium-like peptide 2
CaMKII: Ca^2+^/calmodulin-dependent kinase II
Ca^2+^-free HBSS: Ca^2+^-, Mg^2+^-free Hanks’ balanced salt solution
CRAC: Ca^2+^ release-activated Ca^2+^ channel
CaV1.2: channel subunit of VACC
EF-hand: a calcium-binding motif
EGTA: ethylene glycol-bis(β-aminoethyl ether)-N,N,N’,N’-tetraacetic acid
ER: endoplasmic reticulum
CPA: cyclopiazonic acid
FITC: fluorescein isothiocyanate
HBSS: Hanks balanced salt solution
HRP: horseradish peroxidase
IP_3_: inositol 1,4,5 trisphosphate
IP_3_R: IP_3_ receptor
MLCK: myosin light chain kinase
PI3-K: phosphoinositide 3-kinase
PKC: protein kinases C
PTEN: phosphatase and TENsin homolog deleted on chromosome 10
RRP: rapidly releasable pool
SOCE: store-operated calcium entry
STIM1: stromal-interacting molecule 1
TRPC1: canonical transient receptor potential
Vamp2: vesicle-associated membrane protein 2
VACC: voltage-activated calcium channel

## 10 Acknowledgments

We thank Katie Grzymkowski, Nicole Haessly, Blair Baumle, Jessica Barnett, Robyn Duckworth, and Kaitlyn Neik (Bowling Green State University) for technical assistance. Ms. Lena Scott and Drs. Anita Aperia and Kalaiselvan Krishnan (Karolinska Institutet and Science for Life Laboratory, Sweden) generously provided materials and advice. We are grateful to M. Geusz, D. Giovannucci, M. Model, J. Kozak, and R. Cheney for helpful discussions.

