## Supplementary Material for "How Filopodia Respond to Calcium in the Absence of a Calcium-binding Structural Protein: They Use Rapid Transit"

### 1 Supplementary Figures and Tables

#### 1.1 Supplementary Figures

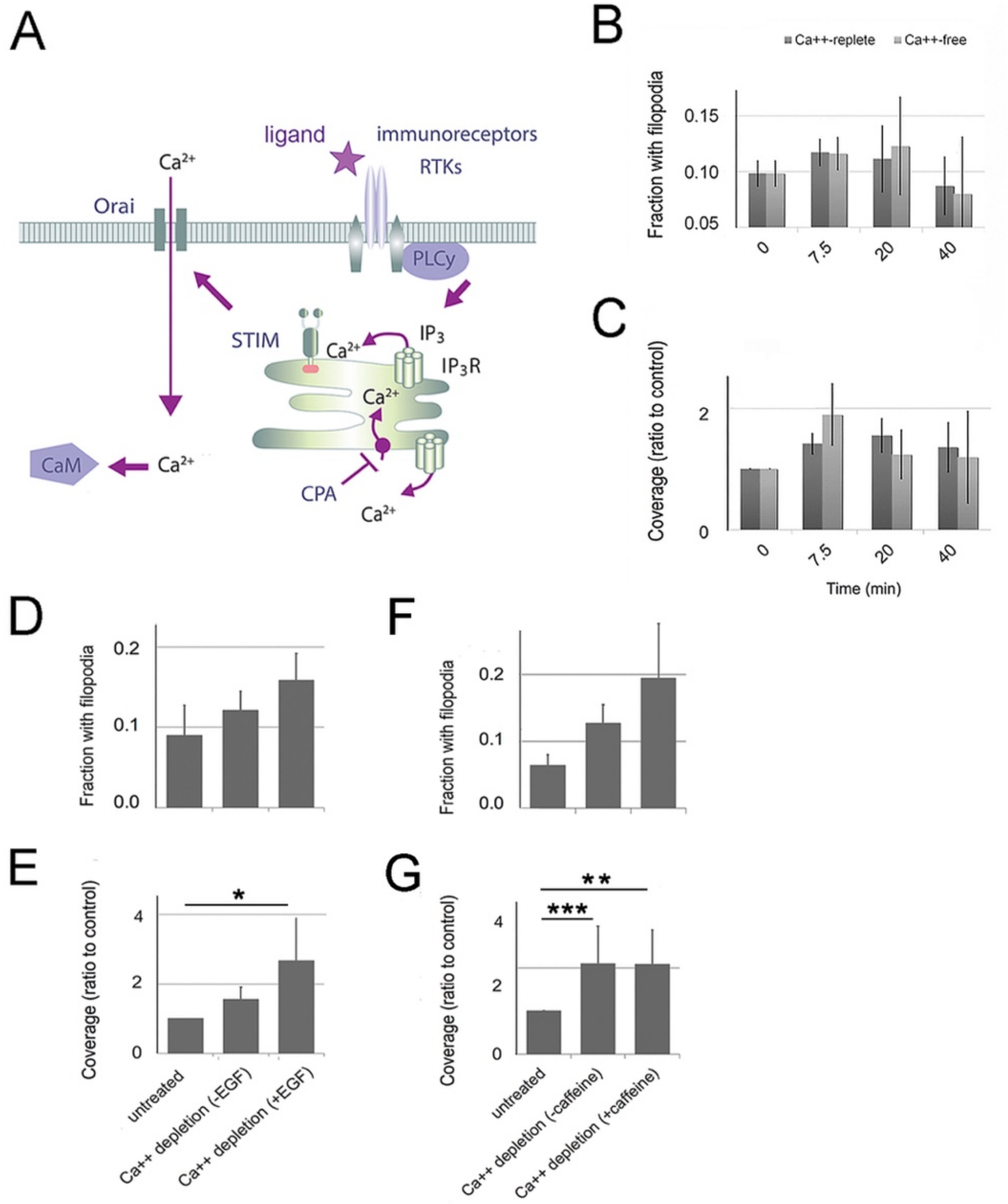


**Supplementary Figure 1.** SOCE initiation leads to changes in filopodia. (A) Canonical pathway of SOCE induction downstream of ligand-receptor interactions (star). Inositol 1,4,5-trisphosphate ($IP_3$)

production is initiated by activation of receptor tyrosine kinases (RTKs) upstream of phospholipase Cγ (PLCγ). Release of $Ca^{2+}$ through the $IP_3$ receptor ($IP_3R$), which is a $Ca^{2+}$ channel, depletes the ER of $Ca^{2+}$ and activates STIM. CPA (cyclopiazonic acid) prevents $Ca^{2+}$ uptake by the SERCA pump (●), thereby preventing its restoration to the ER, so it accumulates in the cytoplasm. (B-C) The culture medium is replaced with HBSS with ($Ca^{2+}$-replete) or without ($Ca^{2+}$-free) $Ca^{2+}$ and filopodia prevalence is analyzed. One-way ANOVA on the experiments had P values from 0.469 to 0.759. (D-E) Cells were exposed for 20 minutes to 5 μM CPA in CMF-HBSS in the presence or absence of 10 ng/ml epidermal growth factor (EGF) and the prevalence of filopodia analyzed after CPA washout with CMF-HBSS and restoration of extracellular $Ca^{2+}$. Prevalence was little affected by $Ca^{2+}$ replenishment after EGF was absent (cf. Figure 1A-B) but was enhanced after EGF was present. (D) ANOVA on the experiment, P=0.163. (E) ANOVA on the experiment, P=0.0051. *Treatments differed at P=0.0055. (F-G) Cells were exposed for 30 minutes to 5 μM CPA in CMF-HBSS in the presence or absence of 150 mM caffeine and the prevalence of filopodia analyzed after CPA washout with CMF-HBSS and restoration of extracellular $Ca^{2+}$. (F) ANOVA on experiment, P=0.539. (G) ANOVA on the experiment, P=0.046. **Treatments differed at P=0.0078. ***Treatments differed at P=0.0055. The fraction of cells with filopodia was slightly changed after restoration of $Ca^{2+}$ in the caffeine-treated samples, but the difference was not significant. Bars represent ± standard error of the mean (S.E.M.).

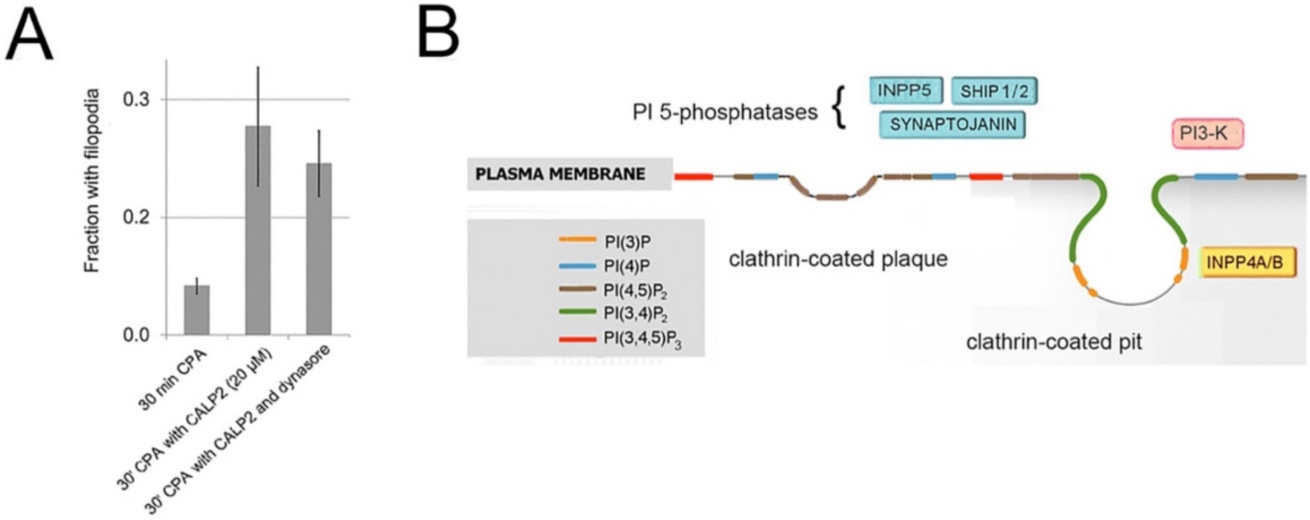


**Supplementary Figure 2.** Contribution of endocytosis to filopodia rescue. (A) Fraction of cells showing filopodia after treatment with CPA alone or with CPA and 20 μM CALP2 in the presence or absence of 45 μM dynasore. ANOVA on the experiment, P=0.00027. (B) Chemotactic cues stimulate the accumulation of phosphatidylinositol 3,4,5-trisphosphate (PI(3,4,5)P3). The substrate of PI3-K, phosphatidylinositol 4,5-bisphosphate (PI(4,5)P2), is present in the plasma membrane especially at clathrin-coated plaques. The product of PI3-K, (PI(3,4,5)P3), is converted to PI(3,4)P2 by PI 5-phosphatases. PI(3,4)P2 participates in endocytic vesicle maturation. PI(3,4)P2 can also be formed by phosphorylation of PI(4)P in an alternate pathway (see for review (Gozzelino et al., 2020). INPP5=PI(4,5)2P 5-phosphatase; SHIP1/2=SH-2 containing inositol 5' polyphosphatase; Synaptojanin=5-phosphatase employing PI(3,4,5)P3 and PI(4,5)P2 as substrates. For details, see references (Schmid, 2014;Stahelin et al., 2014;Bilanges et al., 2019;Dickson and Hille, 2019;Sugiyama et al., 2019).

**Targets of $Ca^{2+}$/calmodulin**

Because the $[Ca^{2+}]_i$ shown in Figure 1C increases continuously, its effect on filopodia dynamics may depend on the channel proteins' binding to calmodulin. It has been reported that calmodulin antagonists potentiate SOCE (Galán et al., 2011), and this could be related to the mechanism of CALP2-mediated filopodia enhancement. Because $Ca^{2+}$-dependent inactivation of both TRPC and Orai was known to be mediated by $Ca^{2+}$/calmodulin (Li et al., 2017), (see for review (Saimi and Kung, 2002;Singh et al., 2002;Bezzerides et al., 2004;Gees et al., 2010;Wang et al., 2020)), their inhibition was considered as a possible mechanism to account for filopodia disappearance. Although calmodulin is thought to be constitutively bound to many TRP channels, *in vitro* studies suggested that the TRPC4 channel only bound $Ca^{2+}$/calmodulin at $Ca^{2+}$ levels exceeding $10^{-5}$ M (Trost et al., 2001). TRPC5 opening is potentiated by $Ca^{2+}$ levels in the same range, however (Blair et al., 2009). Another possible mechanism was $Ca^{2+}$/calmodulin competition at a PI(3,4,5)P3 binding site on TRPC6. $Ca^{2+}$/calmodulin was bound to a C-terminal site, inhibiting the channel, and competition for the site by PI(3,4,5)P3 increased the current through the channel (Kwon et al., 2007). Under this mechanism, wortmannin would inhibit PI(3,4,5)P3 production and TRPC6 activity, which would have negatively affected filopodia prevalence, whereas it had the opposite effect (Figure 2E-F). Two additional mechanisms of CALP2-mediated enhancement were considered. The binding site for $Ca^{2+}$/calmodulin on Orai overlaps its binding site for STIM (Bhardwaj et al., 2020). If the $Ca^{2+}$/calmodulin levels increased, as shown in Figure 1C, STIM1 binding to Orai at the plasma membrane could be blocked inhibiting formation of a STIM1-Orai complex. However, STIM1 became colocalized with Orai in plaques on the plasma membrane during this phase (see **3.1.5 Location of SOCE mediators and aquaporin before and during ER depletion**), suggesting that STIM1-Orai complexes still formed despite a presumed elevation in $Ca^{2+}$/calmodulin. Lastly, all TRPC isoforms shared a C-terminal calmodulin- and $IP_3R$-binding site, where $Ca^{2+}$/calmodulin competes with $IP_3R$ (see for review (Zhu, 2005)). There was a possibility that $Ca^{2+}$/calmodulin dissociated TRPC channels from membrane-bound $IP_3R$ sites. The TRPC1-STIM1 colocalization was unchanged during ER depletion when $[Ca^{2+}]_i$ was maximal, however (see **3.1.5 Location of SOCE mediators and aquaporin before and during ER depletion**). It increased during $Ca^{2+}$ replenishment, when $[Ca^{2+}]_i$ was declining (cf. Figure 1D and Table 2, Supplementary Figure 3C).

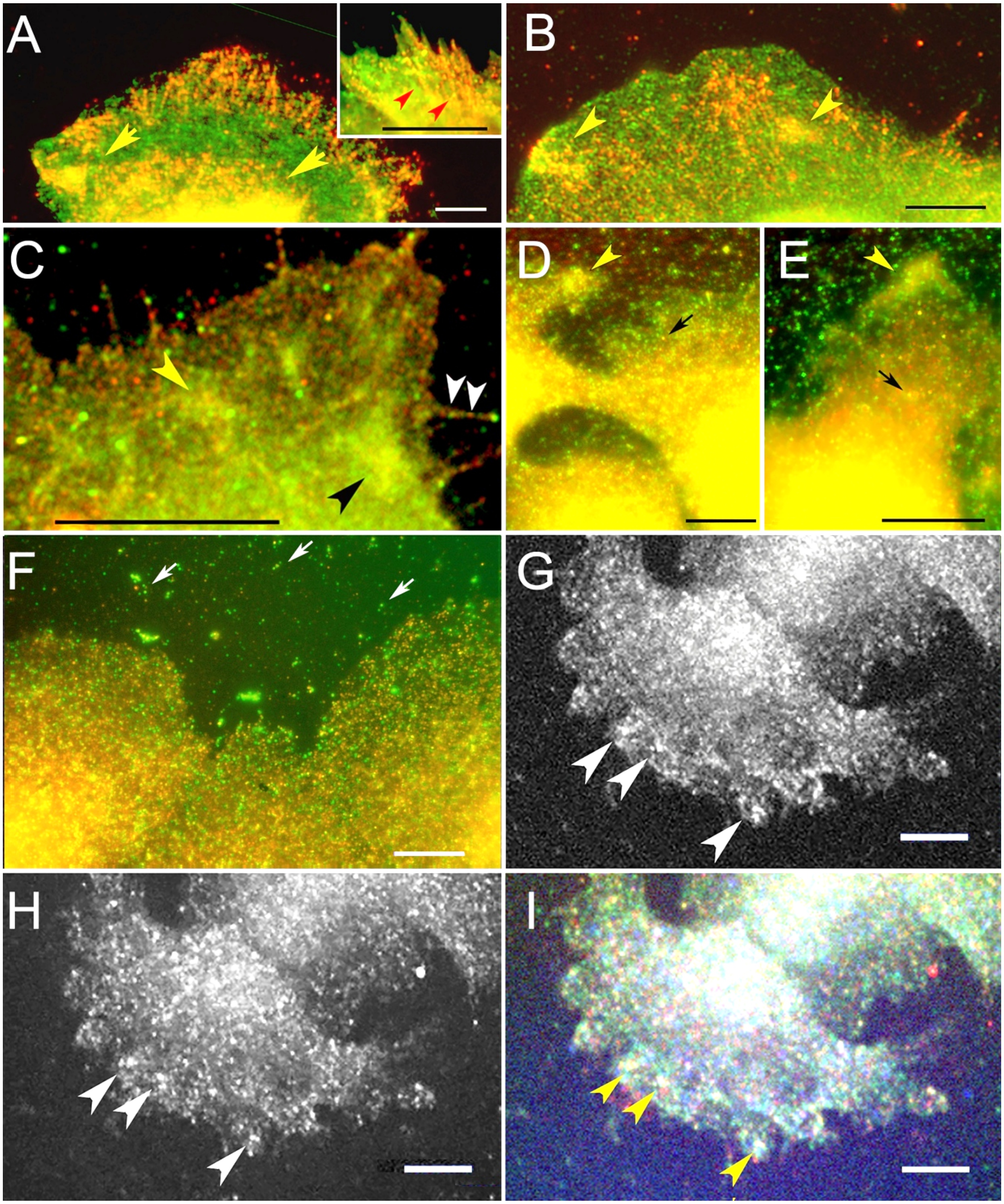


**Supplementary Figure 3.** Redistribution of SOCE mediators during ER depletion and replenishment. (A, B) STIM1 (red) with Orai (green) in amorphous patches (yellow arrowheads). (A) STIM1-Orai1 coincidence (arrowheads) in a cell after ER depletion. Inset: Orai3-STIM1 colocalization showing STIM punctae in linear arrays (red arrowheads), (B) STIM1-Orai1 coincidence (arrowheads) in a cell during $Ca^{2+}$ replenishment, (C) TRPC1 (green) and STIM1 (red) alternating in location on filopodia (white arrowheads) and diffusely distributed on the cell surface (yellow and black arrowheads) during $Ca^{2+}$ replenishment, (D-F) TRPC1 (green) and Vamp2 (red) colocalization. TRPC1 and Vamp2 coincide at elevated portions of the cell (yellow arrowheads).

(D) Vamp2-containing particles at the edge of a cell during $Ca^{2+}$ replenishment. (E) TRPC1-containing particles at the edge of a cell during $Ca^{2+}$ replenishment. (F) Colocalization in untreated cells is mainly in the upper portions of the cell, but the cell edge is distinct. There are a few TRPC1-containing exosomes (white arrows) outside the cells. The largest particles are artifacts from the staining procedure. (G-I) Confocal planes from cells stained with anti-STIM1 (blue), Orai (green), and TRPC1 (red). (G, H) Orai and TRPC1, respectively, in a ventral plane near the lamella (arrowheads), (I) STIM1-Orai colocalization in cyan and TRPC1-Orai colocalization in yellow (arrowheads) in cells during ER depletion. Orai1 and Orai3 showed no difference in the pattern of colocalization with STIM1. Bars =10 µm

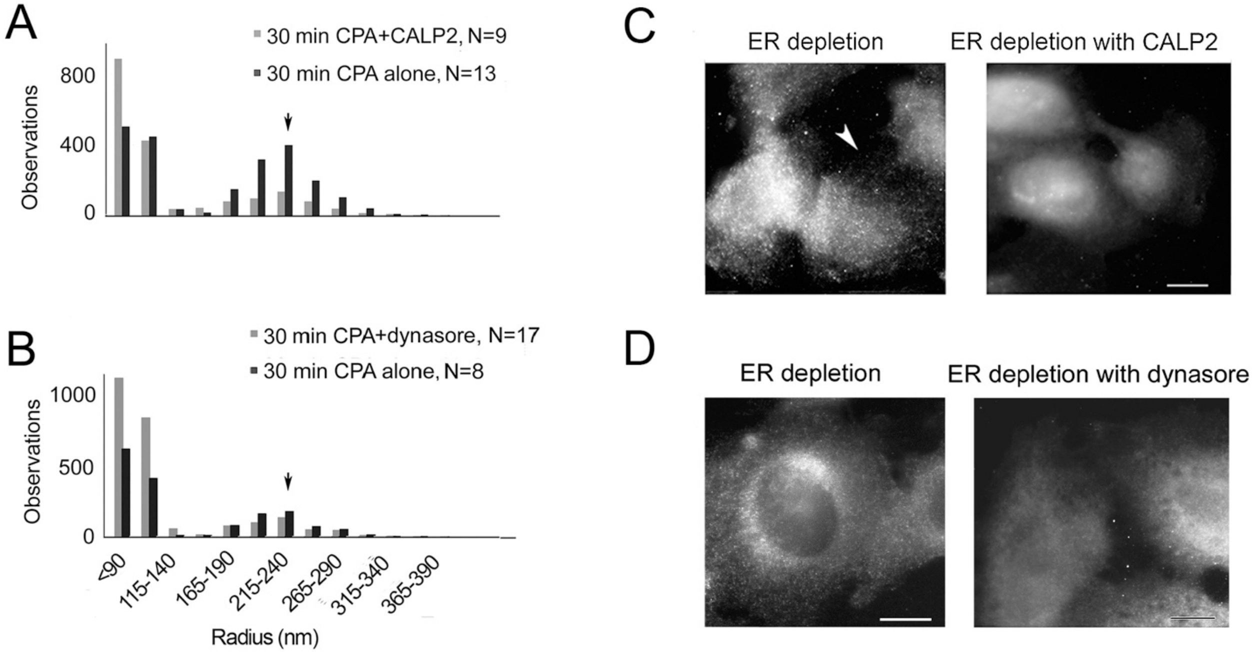


**Supplementary Figure 4.** Results from the presence or absence of agents stimulating filopodia extension during ER depletion. (A-B) TRPC1-bearing loci were localized at the surface of cells that were not permeabilized, and those that had a circular form were segmented for analysis (see **2.11 Analysis of particles' dimensions after staining by indirect immunocytochemistry**). Comparing their distributions in samples during ER depletion showed that the range of radii around 215 nm was represented after CPA alone (cf. Figure 7D), but loci with radii <140 nm predominated in the treated cells. (C-D) Cells were fixed but not permeabilized, and then localization procedures were performed to detect TRPC1 with antibody against the extracellular domain (see **2.7 Immunofluorescence localization, image acquisition, and image processing**). (C) Cells depleted in the presence or absence of 20 µM CALP2. (D) Cells depleted in the presence or absence of 48 µM dynasore. Bars =10 µm

### 1.2 Supplementary Tables

Table S1. Protein pairs with unchanged correlation coefficients in three SOCE phases*

| Phase | STIM1-Orai | STIM1-AQP4 | TRPC1-caveolin | TRPC1-Orai | TRPC-CaV1.2 |
|---|---|---|---|---|---|
| 1 | 0.51 (0.04†, N=10‡) | 0.70 (0.06, N=10) | 0.49 (0.08, N=7) | 0.52 (0.04, N=7) | 0.55 (0.03, N=10) |
| 2 | 0.55 (0.07, N=6) | 0.82 (0.08, N=7) | 0.47 (0.06, N=6) | 0.60 (0.04, N=8) | 0.64 (0.03, N=9) |
| 3 | 0.57 (0.04, N=18) | 0.75 (0.07, N=5) | 0.59 (0.08, N=5) | 0.52 (0.04, N=11) | 0.63 (0.04, N=7) |

*Phase 1 untreated; phase 2 ER depletion; phase 3 $Ca^{2+}$ replenishment
†S.E.M.
‡number of images
